# *Lantana camara* leaf extract-mediated suppression of DNA methyltransferase 1 promotes G0/G1 cell cycle arrest and apoptosis, and impedes migration in MDA-MB-231 triple-negative breast cancer cells

**DOI:** 10.1101/2025.09.26.678791

**Authors:** Sourav Sanyal, Arundhaty Pal, Tapas K Sengupta

## Abstract

**Background:** Triple-negative breast cancer (TNBC) is an aggressive subtype of breast cancer with limited therapeutic options and poor prognosis. DNA methyltransferase 1 (DNMT1) maintains aberrant silencing of tumor suppressor genes, contributing to cancer progression. Our previous work demonstrated that *Lantana camara* leaf extract induces G0/G1 arrest and apoptosis, while inhibiting migration in MDA-MB-231 TNBC cells (Pal et al., 2024).

**Purpose:** This study investigated the role of DNMT1 in mediating the anticancer effects of *Lantana camara* extract.

**Study Design:** An *in vitro* experimental study was performed using MDA-MB-231 TNBC cells.

**Methods:** Cells were treated with the extract, and the expression of DNMT1 was analyzed at the mRNA and protein levels. Expression of tumor suppressor genes was assessed at mRNA level. Functional assays were conducted following DNMT1 overexpression to examine its impact on extract-induced effects. Expression of methyl-CpG binding domain protein 2 (MBD2), histone deacetylase 1 (HDAC1), and histone deacetylase 2 (HDAC2) was evaluated at the mRNA level.

**Results:** The extract significantly downregulated DNMT1, resulting in the reactivation of multiple tumor suppressor genes. Overexpression of DNMT1 counteracted the extract-induced reduction in cell viability, reduced G0/G1 arrest, attenuated apoptosis, and enhanced migratory capacity, while suppressing the mRNA expression of tumor suppressor genes. Additionally, the extract downregulated the mRNA expression of MBD2, HDAC1, and HDAC2, suggesting broader modulation of epigenetic regulators.

**Conclusion:** Suppression of DNMT1 is a central mechanism underlying the anticancer activity of *Lantana camara* extract. These findings highlight its potential as a natural epigenetic modulator and a promising therapeutic candidate for TNBC.

## 1. Introduction

Breast cancer is a major clinical burden, with approximately 2.3 million new cases and 670,000 deaths reported in 2022 (Bray et al., 2024). Breast cancer is a heterogeneous disease comprising several molecular subtypes with distinct therapeutic responses and prognoses. Among these, triple-negative breast cancer (TNBC), characterized by the absence of estrogen receptor, progesterone receptor, and human epidermal growth factor receptor 2 (HER2), represents approximately 15–20% of all breast cancer cases (Lee, 2023). TNBC is highly aggressive, metastatic, and associated with higher rates of recurrence compared to other subtypes (Liang et al., 2020). Current treatment for TNBC mainly relies on chemotherapies due to the lack of defined molecular targets (Yin et al., 2020). However, systemic toxicity and chemoresistance limit therapeutic efficacy, emphasizing the urgent need for novel and more effective treatment strategies.

Epigenetic alterations have emerged as critical drivers of carcinogenesis and therapeutic resistance in breast cancer. Unlike genetic mutations, epigenetic modifications such as DNA methylation, histone modifications, and non-coding RNAs are reversible, making them attractive therapeutic targets (Lu et al., 2020). Aberrant DNA methylation is a characteristic of cancer, leading to the silencing of tumor suppressor genes (Lu et al., 2020). DNA methyltransferases (DNMTs) catalyze the addition of methyl groups to cytosine residues in CpG dinucleotides, maintaining epigenetic gene silencing (Lu et al., 2020). Among them, DNA methyltransferase 1 (DNMT1) is primarily responsible for preserving methylation patterns during DNA replication and remains frequently overexpressed in TNBC (Z. Li et al., 2022; Wong, 2021). Elevated DNMT1 levels are associated with enhanced proliferation, evasion of apoptosis, and increased migratory capacity in TNBC cells (Z. Li et al., 2022). Therefore, DNMT1 has gained attention as a promising target in TNBC.

Several synthetic DNMT inhibitors, such as 5-azacytidine and decitabine, have been approved for clinical use in hematological malignancies (Lu et al., 2020). However, their use in solid tumors like breast cancer is limited due to instability, off-target effects, and systemic toxicity. This has raised growing interest in natural products as alternative epigenetic modulators. Phytochemicals have historically played a central role in the development of anticancer drugs, with notable examples including paclitaxel, vinblastine, and camptothecin (Soumya et al., 2021). A growing body of evidence suggests that phytochemicals can also regulate epigenetic mechanisms, including inhibition of DNMT activity, reversal of aberrant DNA methylation, and reactivation of tumor suppressor genes. Natural compounds such as epigallocatechin gallate, curcumin, and genistein have been shown to exert DNMT inhibitory activity, thereby restoring normal gene expression patterns and suppressing tumor progression (Aanniz et al., 2024; Saldívar-González et al., 2018). These findings highlight the potential of phytochemicals as epigenetic therapies for aggressive cancers such as TNBC.

It was reported previously from our laboratory that *Lantana camara* leaf ethanolic extract (LCLEE) exerts its cytotoxic effects by inducing G0/G1 cell cycle arrest and apoptosis, and also inhibits the migration of MDA-MB-231 TNBC cells (Pal et al., 2024). However, the molecular mechanism exerting these effects remains unexplored.

In this study, the effects of LCLEE on MDA-MB-231 cells are examined, with a focus on its role in regulating the expression of DNMT1. We investigated whether LCLEE-mediated suppression of DNMT1 was associated with alterations in key cellular processes such as cell cycle progression, apoptosis, and migration using the MDA-MB-231 cell line as a representative model of TNBC.

## 2. Materials and methods

### 2.1. Cell culture

MDA-MB-231 cell line was procured from the National Centre for Cell Science (NCCS), Pune, India. The previously described technique was followed to culture the cells (Pal et al., 2024).

### 2.2. *Lantana camara* leaf ethanolic extract (LCLEE) preparation

LCLEE preparation and treatment protocol are the same as those reported elsewhere (Pal et al., 2024).

### 2.3. RNA isolation, cDNA synthesis, and reverse transcription quantitative real-time polymerase chain reaction (RT-qPCR)

MDA-MB-231 cells were cultured in 60 mm dishes (0.4 x 10^6^ cells/dish) to 70-80% confluency. The cells were then treated with vehicle (0.8% ethanol) and concentrations of LCLEE at 40 µg/mL, 60 µg/mL, 80 µg/mL, 120 µg/mL, and 180 µg/mL for 24 hours. Total RNA was isolated from vehicle-treated and LCLEE-treated MDA-MB-231 cells using RNAiso Plus (Takara, Japan) according to the manufacturer’s instructions.

The RNA samples were further used for the synthesis of cDNA using the PrimeScript 1st strand cDNA synthesis kit (Takara, Japan) according to the manufacturer’s protocol. The detailed protocol was reported previously (Pal et al., 2024).

Real-time PCR was performed using TB Green Premix Ex Taq (Tli RNaseH Plus) kit (Takara, Japan) according to the manufacturer’s instructions. The detailed protocol was described previously (Pal et al., 2024). Primers of *18S*, *DNMT1*, *CDKN1A*, *PTEN*, *FOXO3a*, *Claudin-1*, *MBD2*, *HDAC1*, and *HDAC2* were used for a gene expression study (Supplementary Table S1). The gene expression was normalized against the *18S* rRNA housekeeping gene, and the data were analysed using the 2^-ΔΔCt^ method and plotted as fold change versus vehicle control (Livak and Schmittgen, 2001).

### 2.4. Western Blotting

MDA-MB-231 cells were grown in 60 mm dishes (0.4 x 10^6^ cells/dish) till 70-80% confluency, followed by treatment with vehicle (0.8% ethanol) and concentrations of LCLEE at 40 µg/mL, 60 µg/mL, 120 µg/mL, and 180 µg/mL for 24 hours, along with untreated control cells. The cells were harvested by trypsinization and washed using phosphate-buffered saline (PBS). Lysis of the cells was performed using Radioimmunoprecipitation assay (RIPA) buffer (HiMedia, India), supplemented with protease inhibitor cocktail (Sigma-Aldrich, USA), with frequent agitation for 30 minutes on ice. A protein lysate was then obtained as supernatant by centrifugation at 12,000 rpm for 20 minutes at 4 °C. Bradford assay was performed to measure the protein concentration in the cell lysate. An equal amount of total protein (50 μg) from all treatment conditions was separated by electrophoresis on 8% polyacrylamide gel and then electro-transferred onto a polyvinylidene fluoride (PVDF) membrane. The membrane was blocked with 5% (w/v) bovine serum albumin (BSA) fraction V (Amresco, USA) in TBST. The membrane was incubated overnight at 4°C with the respective primary antibody, followed by incubation with horseradish peroxidase (HRP)-conjugated secondary antibody. Finally, the protein bands were detected using a chemiluminescent substrate (SuperSignal West Pico, Thermo Fisher Scientific, USA) and the G:BOX Chemi XRQ imaging system (Syngene). The NIH-ImageJ 1.53k software was used to quantify the intensity of the protein bands, which were then normalized using respective GAPDH (loading control) protein bands. The details of the antibodies are listed in Supplementary Table S2.

### 2.5. Generation of polyclonal stable cell lines

To generate stable MDA-MB-231 cell lines, cells were transfected with either pcDNA3.1(+) empty vector encoding a neomycin resistance gene (neoR) or pcDNA3.1(+)-DNMT1 construct encoding both the transgene (DNMT1) and a neomycin resistance gene (neoR). Details of the plasmid and cloning procedure are provided in the supplementary section (Supplementary Figure S1). MDA-MB-231 cells were seeded at a density that allowed them to reach ∼80% confluence at the time of transfection. Transfection was performed using Lipofectamine^TM^ 2000 Transfection Reagent (ThermoFisher Scientific, USA) following the manufacturer’s instructions. 48 hours post-transfection, the culture medium was replaced with fresh complete medium supplemented with G-418 sulphate (Geneticin) (SRL, India).

For selection, transfected cells were cultured in complete medium containing 700 µg/mL G-418 sulphate (a predetermined concentration). Selection was continued for 14 days, with media changes every 3 days, until non-resistant cells were eliminated and resistant colonies emerged. Surviving cells were expanded in the presence of G-418 to generate a polyclonal stable cell population.

Stable integration and expression of DNMT1 were verified at both the mRNA and protein levels. Here, the polyclonal stable cell line containing the empty vector is termed MDA-MB-231/pcDNA3.1(+), whereas the polyclonal stable cell line containing the pcDNA3.1(+)-DNMT1 construct is termed MDA-MB-231/pcDNA3.1(+)-DNMT1. For long-term maintenance, cells were cultured in G-418 sulphate-containing medium at a maintenance concentration (200 µg/mL).

### 2.6. Cell viability assay

The effect of LCLEE on the viability of MDA-MB-231/pcDNA3.1(+) and MDA-MB-231/pcDNA3.1(+)-DNMT1 cells was evaluated by the 3-(4,5-dimethylthiazol-2-yl)-2,5-diphenyltetrazolium bromide (MTT) colorimetric assay following the same protocol reported previously (Pal et al., 2024).

### 2.7. Cell cycle phase distribution assay

MDA-MB-231/pcDNA3.1(+) and MDA-MB-231/pcDNA3.1(+)-DNMT1 cells were grown in 35 mm cell culture dishes (0.2 x 10^6^ cells/dish) till 70-80% confluency. Cells were then treated with vehicle (0.8% ethanol) and concentrations of LCLEE at 60 µg/mL, 80 µg/mL, 100 µg/mL, 120 µg/mL, 150 µg/mL, and 180 µg/mL for 24 hours, along with untreated control cells. The rest of the protocol is the same as published elsewhere (Pal et al., 2024).

### 2.8. Annexin V-Fluorescein isothiocyanate (FITC)/Propidium iodide (PI) double staining

The detection of apoptosis post-treatment with the extract was performed using the Annexin V-FITC Apoptosis detection kit (BD Biosciences, USA) according to the manufacturer’s protocol. Briefly, MDA-MB-231/pcDNA3.1(+) and MDA-MB-231/pcDNA3.1(+)-DNMT1 cells were grown in 35 mm cell culture dishes (0.2 x 10^6^ cells/dish) till 70-80% confluency. Cells were then treated with vehicle (0.8% ethanol) and concentrations of LCLEE at 60 µg/mL, 120 µg/mL, 180 µg/mL, 240 µg/mL, and 300 µg/mL for 24 hours, along with untreated control cells. The remainder of the protocol is identical to that described elsewhere (Pal et al., 2024).

### 2.9. Wound healing assay

Wound healing assay was performed following the protocol discussed previously (Pal et al., 2024). Briefly, MDA-MB-231/pcDNA3.1(+) and MDA-MB-231/pcDNA3.1(+)-DNMT1 cells were cultured in 35 mm dishes (0.3 x 10^6^ cells/dish) and grown till the formation of a uniform monolayer. A wound was made in each dish using a sterile 0.2-10 μL micro pipette tip, and the consumed media was discarded from the culture dish. Then, the monolayer of cells was washed once with 1X PBS. Cells were then treated with concentrations of 40 μg/mL, 60 μg/mL, 80 μg/mL, 100 μg/mL, and 120 μg/mL LCLEE, along with vehicle-treated and untreated control cells. 0.5 μg/mL cycloheximide was added in every treatment condition to inhibit cell proliferation. Images of the cells were captured at 0 hours and 24 hours post-treatment using EVOS XL Core Cell Imaging System (Invitrogen, ThermoFisher Scientific, USA) at 10X magnification. The images were analysed, and the area of the wounds was calculated by ImageJ 1.53k (ImageJ software, NIH, USA) using the following formula:

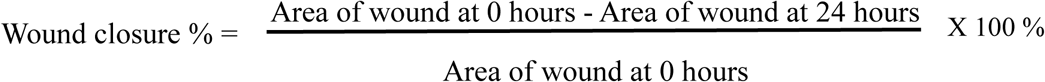

### 2.10. Statistical analysis

All the experiments were conducted at least three times. Data were plotted using GraphPad Prism 8.0.1 (GraphPad software, CA, USA) and represented as mean ± SEM (Standard Error of Mean). Statistical analysis was done by performing a parametric Student’s t-test. P<0.05 was considered to indicate a statistically significant difference.

## 3. Results

### 3.1. LCLEE downregulated the mRNA and protein expression of DNMT1

DNMT1 plays a significant role in promoting the development and aggressive behavior of TNBC. DNMT1 remains overexpressed in TNBC, and its increased expression is associated with poorer survival outcomes (Z. Li et al., 2022; Wong, 2021). It contributes to tumorigenesis by silencing tumor suppressor genes through hypermethylation of their promoter regions (Wong, 2021; Yu et al., 2019). Given the oncogenic role and elevated expression of DNMT1 in TNBC, we examined whether LCLEE modulates DNMT1 expression in MDA-MB-231 TNBC cells. DNMT1 expression was analyzed at both the mRNA and protein levels following 24 hours of LCLEE treatment. The mRNA expression of *DNMT1* decreased dose-dependently in MDA-MB-231 cells treated with LCLEE compared to vehicle control. *DNMT1* mRNA expression (Figure 1A) was found to be significantly downregulated to 0.55 ± 0.02, 0.36 ± 0.01, 0.30 ± 0.03, and 0.26 ± 0.04-fold at 40 µg/mL, 60 µg/mL, 120 µg/mL, and 180 µg/mL concentrations of LCLEE, respectively. When the protein expression of DNMT1 was checked under the same treatment conditions, it was found that the protein level of DNMT1 (Figure 1B and 1C) was decreased to 0.40 ± 0.10 and 0.53 ± 0.21-fold at 40 µg/mL and 60 µg/mL concentrations of LCLEE, respectively, with very little or no expression at 120 µg/mL and 180 µg/mL concentrations.

**Figure 1.**
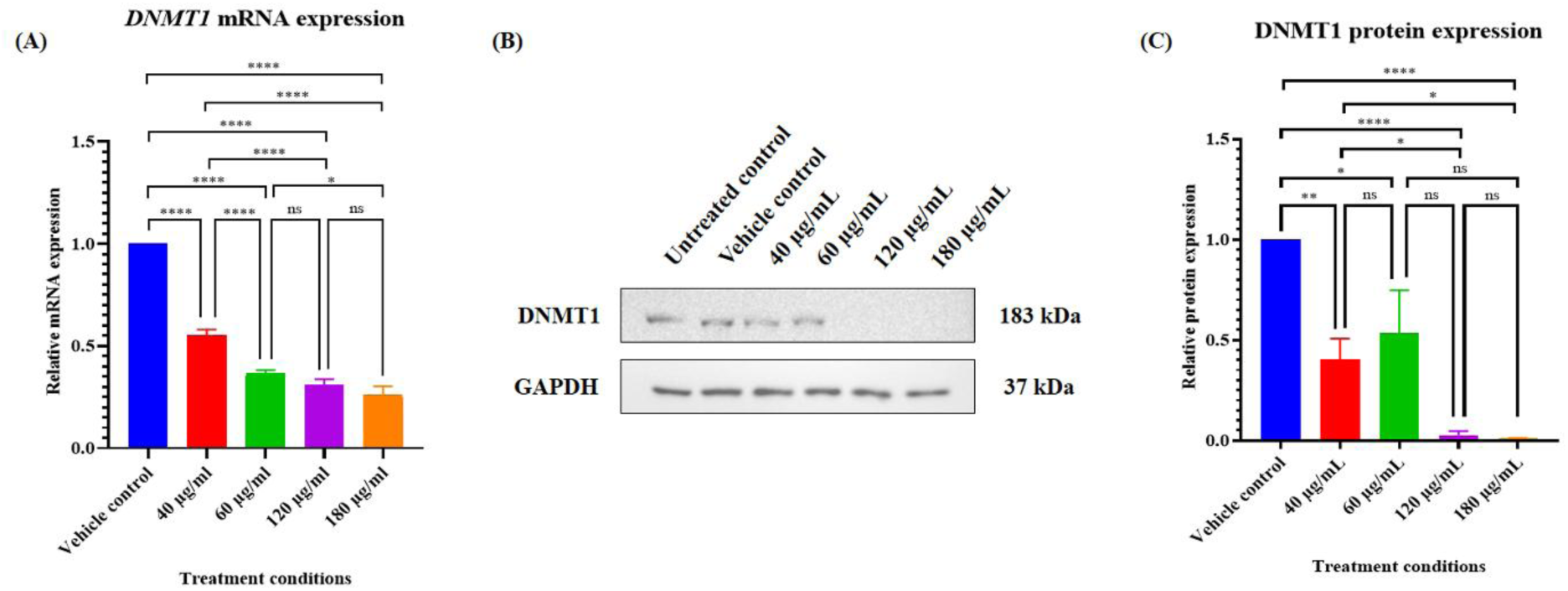
mRNA and protein expression of DNA methyltransferase 1 were altered in MDA-MB-231 cells compared to vehicle control when treated with various doses of *Lantana camara* leaf ethanolic extract for 24 hours. **(A)** relative mRNA expression of *DNMT1*, **(B)** Representative western blot images, and **(C)** normalized graph of DNMT1 protein expression. Data is represented as mean ± SEM of three independent biological replicates, where * is *P*≤0.05, ** is *P*≤0.01, **** is *P*≤0.0001, and ns signifies non-significant.

### 3.2. LCLEE upregulated the mRNA expression of multiple tumor suppressor genes

Several studies have reported that overexpression of DNMT1 silences tumor suppressor genes that are involved in cell proliferation, apoptosis, and migration, such as *CDKN1A* (Al-Kharashi et al., 2017; Mirza et al., 2013), *PTEN* (Liu et al., 2021; Stefanska et al., 2012), *FOXO3a* (Liu et al., 2019), and *Claudin-1* (Chiang et al., 2019; Zhu et al., 2024). It was observed that LCLEE downregulated the mRNA and protein expression of DNMT1. Since DNMT1 primarily regulates gene expression through DNA methylation at promoter regions, its effects are mostly reflected at the transcriptional level. Therefore, the mRNA expression of selected tumor suppressor genes (*CDKN1A*, *PTEN*, *FOXO3a*, and *Claudin-1*), which are known to be silenced by DNMT1 overexpression, was examined in MDA-MB-231 cells treated with various concentrations of LCLEE, along with a vehicle control. Compared to vehicle control, the transcript level of *CDKN1A* (Figure 2A) increased to 3.56 ± 0.31, 4.14 ± 0.21, 7.87 ± 2.08, and 9.09 ± 2.00-fold when treated with 40 µg/mL, 60 µg/mL, 120 µg/mL, and 180 µg/mL concentrations of LCLEE, respectively. *PTEN* mRNA expression (Figure 2B) was found to be significantly upregulated to 1.32 ± 0.09, 1.46 ± 0.10, 1.64 ± 0.26, and 1.65 ± 0.15-fold when MDA-MB-231 cells were treated with LCLEE at 40 µg/mL, 60 µg/mL, 120 µg/mL, and 180 µg/mL, respectively. Similarly, *FOXO3a* mRNA expression (Figure 2C) was found to be significantly upregulated to 1.27 ± 0.10, 1.51 ± 0.21, and 2.23 ± 0.12-fold when MDA-MB-231 cells were treated with 60 µg/mL, 120 µg/mL, and 180 µg/mL concentrations of LCLEE, respectively; however, no significant change in the mRNA expression was observed at 40 µg/mL concentration. As Claudin-1 is primarily involved in cell migration, its mRNA expression was assessed at lower concentrations. The transcript level of *Claudin-1* (Figure 2D) was found to be significantly upregulated to 2.93 ± 0.68, 5.06 ± 1.10, and 6.42 ± 1.79-fold when treated with 40 µg/mL, 60 µg/mL, and 80 µg/mL concentrations of LCLEE, respectively, compared to vehicle control.

**Figure 2.**
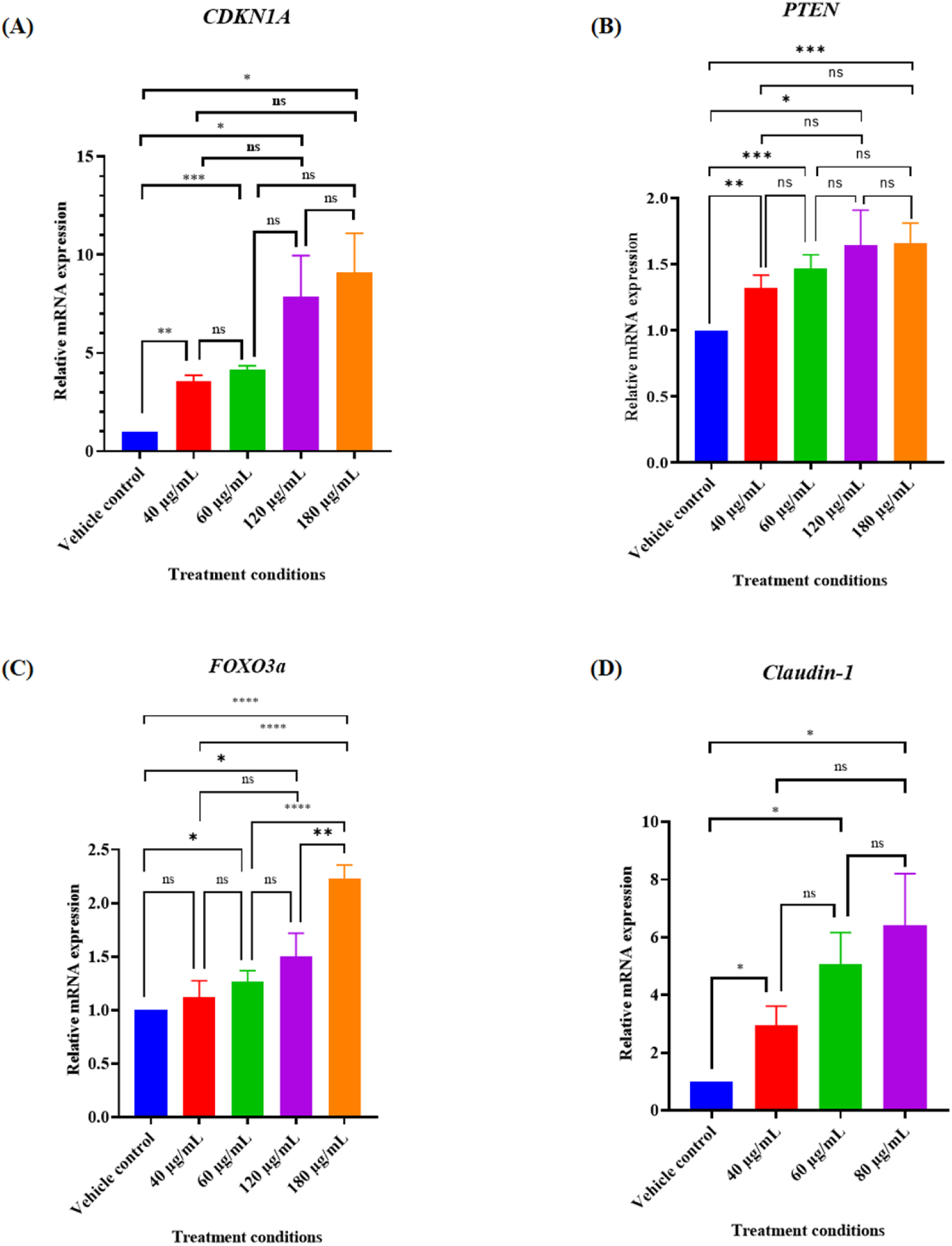
mRNA expression levels of tumour suppressor genes were upregulated in MDA-MB-231 cells compared to vehicle control when treated with various doses of *Lantana camara* leaf ethanolic extract for 24 hours. Relative expression of **(A)** *CDKN1A*, **(B)** *PTEN*, **(C)** *FOXO3a*, and **(D)** *Claudin-1*. Data is represented as mean ± SEM of three independent biological replicates, where * is P≤0.05, ** is P≤0.01, *** is P≤0.001, **** is P≤0.0001, and ns signifies non-significant.

### 3.3. Screening of DNMT1 expression in polyclonal stable cell lines

To place the findings on DNMT1 suppression within a broader functional context, the existing literature was examined to find the consequences of DNMT1 loss in cancer cells. Pan and colleagues demonstrated that the knockdown of DNMT1 in MDA-MB-231 cells resulted in reduced viability and proliferation, accompanied by the induction of apoptosis, highlighting the critical role of DNMT1 in cancer cell proliferation and survival (Pan et al., 2022). Another study revealed that suppression of DNMT1 significantly impairs the migratory ability of MDA-MB-231 cells (H. Li et al., 2022). Taken together, these studies indicate that DNMT1 acts as a central regulator of multiple malignant phenotypes, including proliferation, survival, and migration.

In our previous study, we observed that LCLEE treatment reduced cell viability, induced G0/G1 cell cycle arrest and apoptosis, and inhibited the migration of MDA-MB-231 TNBC cells (Pal et al., 2024). In the present study, we observed that LCLEE downregulated DNMT1 and upregulated the expression of multiple tumor suppressor genes involved in cell cycle regulation, apoptosis, and migration. This led us to examine whether DNMT1 downregulation is mechanistically responsible for the LCLEE-mediated effects. To address this, DNMT1 was ectopically overexpressed in MDA-MB-231 cells to determine whether restoring DNMT1 could attenuate or reverse LCLEE-induced effects on proliferation, survival, and migration. This approach directly tested the causal role of DNMT1 downregulation in mediating the anticancer activities of LCLEE.

To validate the successful generation of DNMT1-overexpressing polyclonal stable cell lines, DNMT1 expression was evaluated at both the mRNA and protein levels. As shown in Figure 3A, MDA-MB-231 cells stably transfected with the pcDNA3.1(+)-DNMT1 construct exhibited a significant increase in DNMT1 transcript levels compared to cells transfected with the empty pcDNA3.1(+) vector. Consistent with the transcript data, immunoblot analysis also confirmed a significant increase in DNMT1 protein expression in MDA-MB-231 cells stably transfected with the pcDNA3.1(+)-DNMT1 construct relative to empty vector control (Figure 3B). These results confirm the generation of polyclonal stable cell lines with sustained ectopic overexpression of DNMT1 and validate their suitability for subsequent functional and mechanistic analyses.

**Figure 3.**
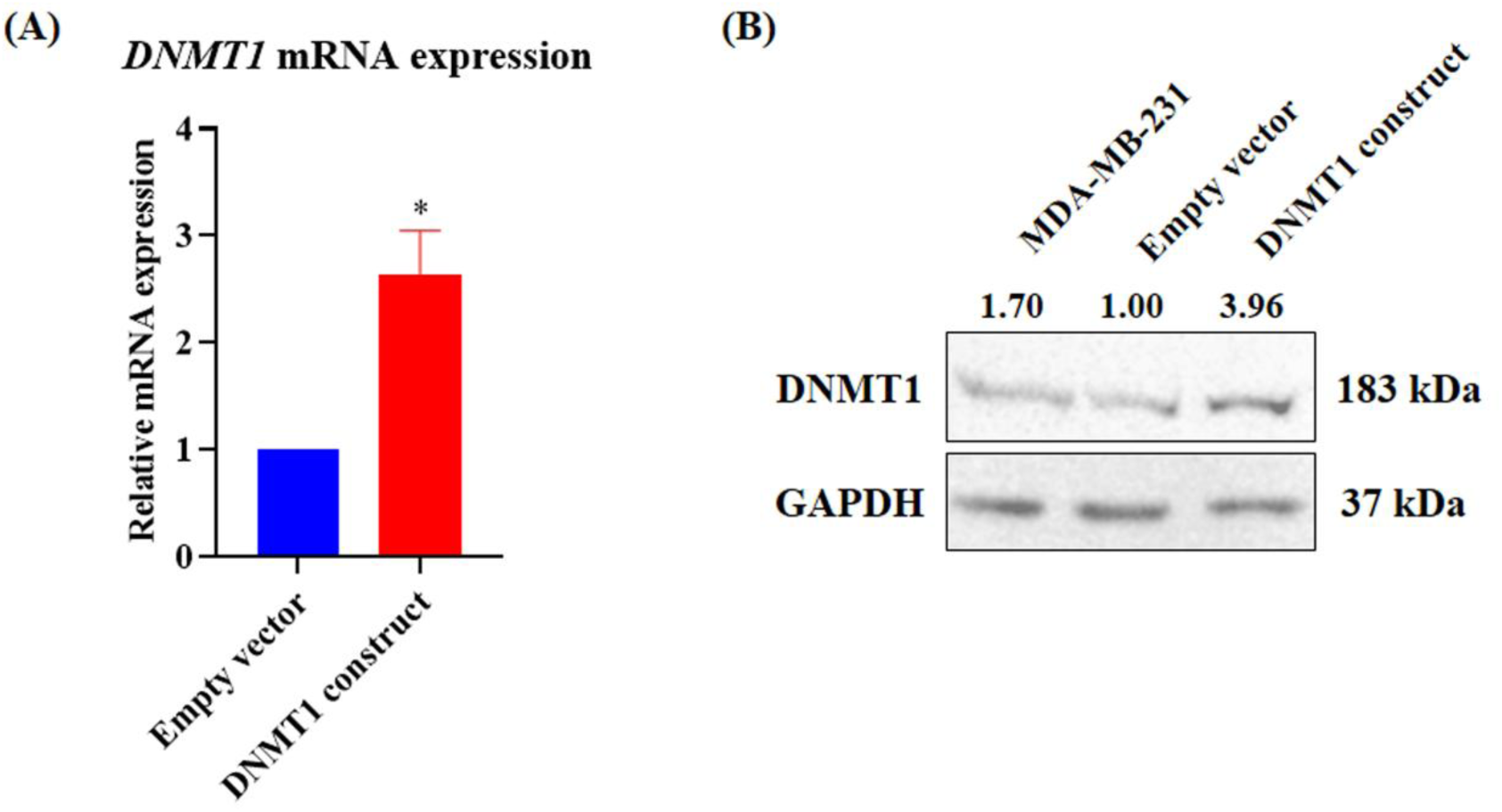
Screening and validation of DNMT1 overexpression in polyclonal stable MDA-MB-231 cell lines. **(A)** relative mRNA expression of *DNMT1*, **(B)** Representative western blot images with normalized values of DNMT1 protein expression. Data is represented as mean ± SEM of three independent biological replicates, where * signifies *P*≤0.05.

### 3.4. Ectopic overexpression of DNMT1 rescued cell viability

To examine whether DNMT1 overexpression influences the cytotoxic response to LCLEE, cell viability was evaluated in LCLEE-treated polyclonal stable MDA-MB-231 cell lines expressing either the empty pcDNA3.1(+) vector or pcDNA3.1(+)-DNMT1 construct using the MTT assay. As shown in Figure 4, LCLEE treatment resulted in a clear dose-dependent reduction in cell viability following 24 hours of exposure in empty-vector cells. In contrast, DNMT1-overexpressing cells exhibited a significantly attenuated reduction in viability at corresponding LCLEE concentrations. Notably, higher concentrations of LCLEE were required to achieve comparable levels of growth inhibition in DNMT1-overexpressing cells relative to empty-vector control. These findings indicate that ectopic DNMT1 overexpression partially rescues MDA-MB-231 cells from LCLEE-induced growth suppression, suggesting a functional role of DNMT1 in mediating the cytotoxic response to LCLEE.

**Figure 4.**
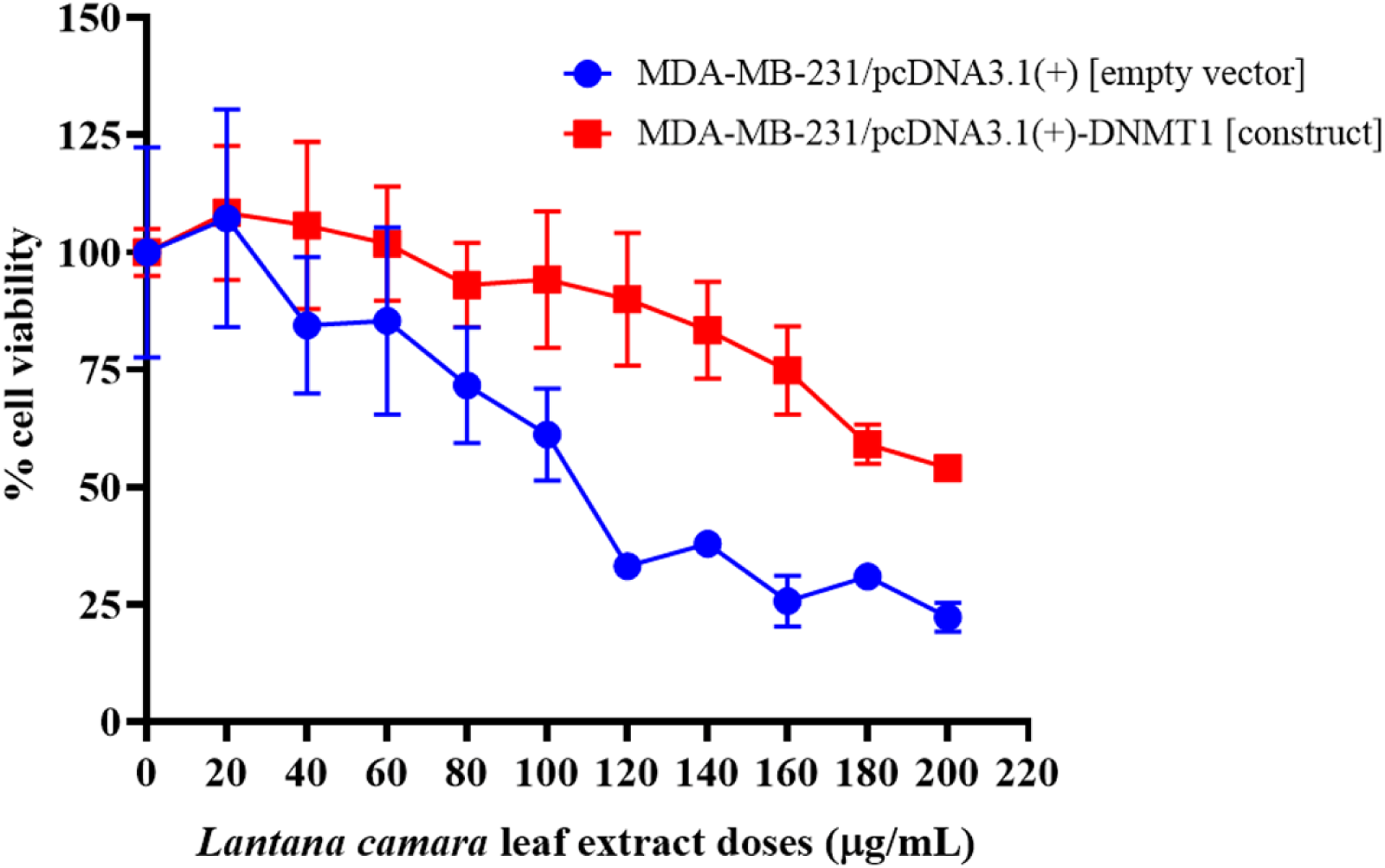
Effect of LCLEE on cell viability in empty-vector and DNMT1-overexpressing MDA-MB-231 cells. Polyclonal stable MDA-MB-231 cells expressing empty pcDNA3.1(+) vector or pcDNA3.1(+)-DNMT1 construct were treated with increasing concentrations of LCLEE for 24 hours, and cell viability was assessed by MTT assay. Data are presented as percentage cell viability relative to vehicle control and expressed as mean ± SD.

### 3.5. DNMT1 overexpression attenuated LCLEE-mediated G0/G1 arrest

To further investigate whether the downregulation of DNMT1 is responsible for the anti-proliferative effect of the extract, cell cycle phase distribution was analysed in empty vector and DNMT1-overexpressing MDA-MB-231 cells to compare the effect of LCLEE. Figure 5A and 5C are representative images of the distribution of MDA-MB-231/pcDNA3.1(+) and MDA-MB-231/pcDNA3.1(+)-DNMT1 cells in distinct cell cycle phases when treated with a range of concentrations of the extract from 60 μg/mL to 180 μg/mL for 24 hours, along with vehicle-treated and untreated controls. Analyses of the results (Figure 5B) revealed that G0/G1 population of MDA-MB-231/pcDNA3.1(+) cells increased from 60.967 ± 0.878% (vehicle control) to 65.917 ± 1.049%, 68.833 ± 1.160%, 72.467 ± 0.495%, 74.517 ± 0.588%, 72.817 ± 1.131%, and 68.700 ± 1.479% when treated with 60 μg/mL, 80 μg/mL, 100 μg/mL, 120 μg/mL, 150 μg/mL, and 180 μg/mL concentrations of LCLEE, respectively. Whereas, Figure 5D revealed that the G0/G1 population of MDA-MB-231/pcDNA3.1(+)-DNMT1 cells increased from 66.917 ± 1.097% (vehicle control) to 69.133 ± 0.304%, 71.083 ± 0.776%, 72.617 ± 0.516%, 73.400 ± 0.842%, 73.133 ± 0.638%, and 69.167 ± 0.429% when treated with 60 μg/mL, 80 μg/mL, 100 μg/mL, 120 μg/mL, 150 μg/mL, and 180 μg/mL concentrations of LCLEE, respectively. This finding indicated that treatment with the extract induced G0/G1 arrest in both MDA-MB-231/pcDNA3.1(+) and MDA-MB-231/pcDNA3.1(+)-DNMT1 cells. However, the increase in G0/G1 cell population at the 60 μg/mL concentration of the extract is non-significant in the case of MDA-MB-231/pcDNA3.1(+)-DNMT1 cells compared to the significant increase in MDA-MB-231/pcDNA3.1(+) cells. It is also noteworthy that the extent to which the G0/G1 cell population increased is less in MDA-MB-231/pcDNA3.1(+)-DNMT1 cells compared to MDA-MB-231/pcDNA3.1(+) cells. The observed increase in the G0/G1 population in MDA-MB-231/pcDNA3.1(+) and MDA-MB-231/pcDNA3.1(+)-DNMT1 was associated with a significant decrease in the S-phase population. However, no significant change was observed in the G2/M cell population in MDA-MB-231/pcDNA3.1(+)-DNMT1 cells compared to the MDA-MB-231/pcDNA3.1(+) cells, where G2/M cell population decreased significantly from 60 μg/mL to 150 μg/mL concentrations of LCLEE. A comparative table of cell cycle phase distribution at different treatment conditions is provided in Supplementary Table S4. Taken together, the results established that the ectopic overexpression of DNMT1 can attenuate LCLEE-induced G0/G1 arrest.

**Figure 5.**
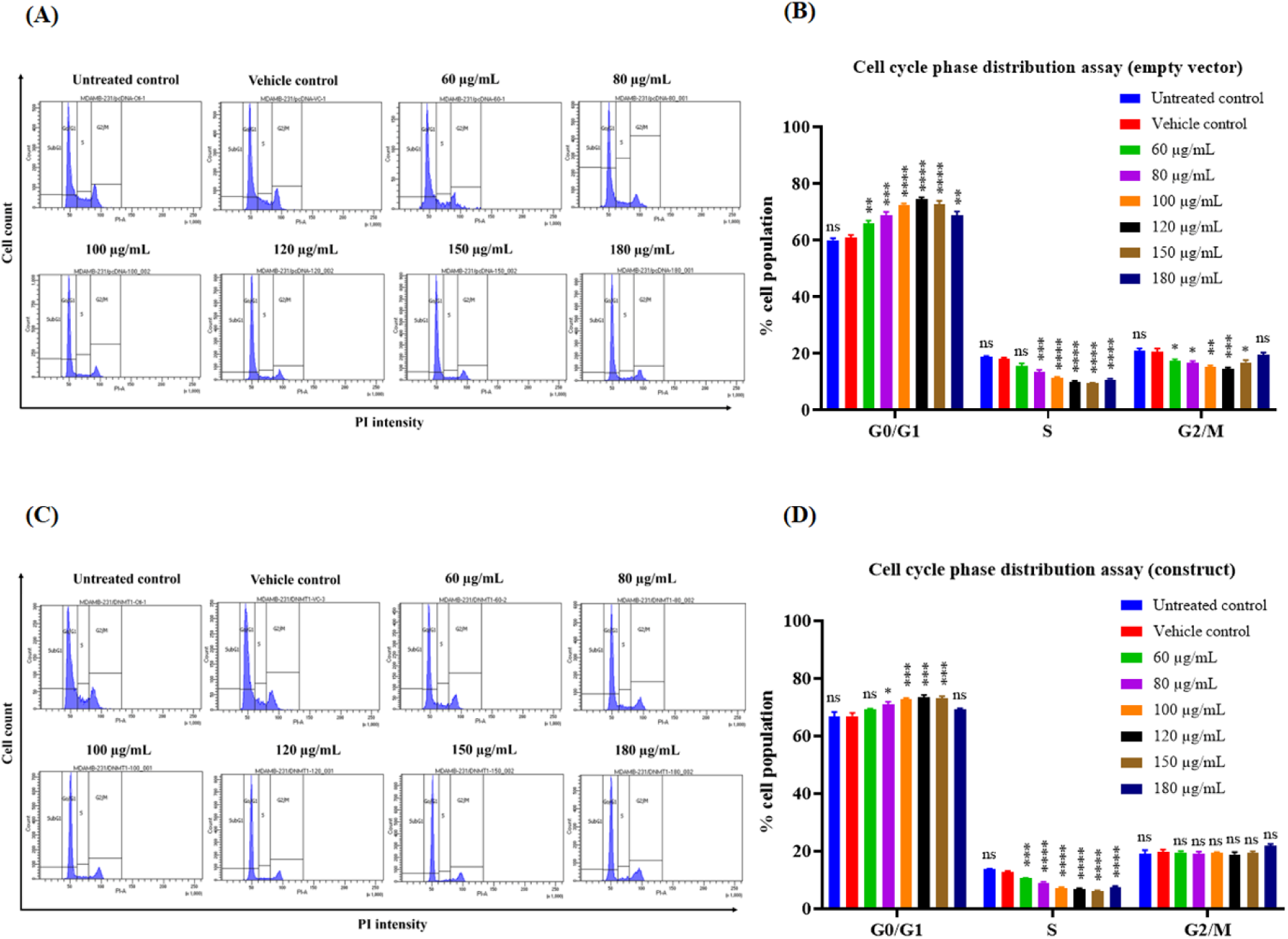
Representative images of cell cycle phase distribution of **(A)** MDA-MB-231/pcDNA3.1(+) cells and **(C)** MDA-MB-231/pcDNA3.1(+)-DNMT1 cells following 24 hours of treatment with the leaf extract at different doses. Bar diagrams showing the percentage of **(B)** MDA-MB-231/pcDNA3.1(+) cells and **(D)** MDA-MB-231/pcDNA3.1(+)-DNMT1 cells in different cell cycle phases (G0/G1, S, and G2/M) following 24 hours of treatment with the extract at various doses. Data are means of three independent experiments and presented as means ± SEM, where * is *P*≤0.05, ** is *P*≤0.01, *** is *P*≤0.001, **** is *P*≤0.0001, and ns signifies non-significant.

### 3.6. Ectopic DNMT1 overexpression countervailed LCLEE-induced apoptosis

To evaluate whether the overexpression of DNMT1 affects the pro-apoptotic activity of the extract, Annexin V-FITC/PI double staining was performed in MDA-MB-231/pcDNA3.1(+) and MDA-MB-231/pcDNA3.1(+)-DNMT1 cells. Figure 6A and 6C are representative images of the scatter plots showing percentage live/viable (Q3), early apoptotic (Q4), late apoptotic (Q2), and non-apoptotic (Q1) populations of MDA-MB-231/pcDNA3.1(+) and MDA-MB-231/pcDNA3.1(+)-DNMT1 cells when treated for 24 hours with 60 μg/mL, 120 μg/mL, 180 μg/mL, 240 μg/mL, and 300 μg/mL concentrations of LCLEE, along with vehicle-treated and untreated controls. Analysis of the results (Figure 6B) revealed that late apoptotic cell population of MDA-MB-231/pcDNA3.1(+) cells increased from 6.275 ± 0.557% (vehicle control) to 10.70 ± 1.901%, 12.975 ± 2.478%, 17.85 ± 4.777%, 18.175 ± 4.826%, and 28.275 ± 7.393%; non-apoptotic cell population increased from 1.90 ± 0.319% (vehicle control) to 4.075 ± 0.68%, 4.475 ± 0.829%, 7.275 ± 1.736%, 18.025 ± 5.693%, and 37.400 ± 12.906% when treated with 60 μg/mL, 120 μg/mL, 180 μg/mL, 240 μg/mL, and 300 μg/mL concentrations of LCLEE, respectively. This finding implied that treatment with LCLEE increased late and non-apoptotic populations of MDA-MB-231/pcDNA3.1(+) cells, and the observed increase was accompanied by a significant concomitant decrease in live or viable cell population, without any change in the early apoptotic cell population compared to the vehicle control. Whereas, analysis of the results (Figure 6D) revealed that the late apoptotic cell population of MDA-MB-231/pcDNA3.1(+)-DNMT1 cells increased significantly from 6.20 ± 0.484% (vehicle control) to 10.083 ± 0.712 and 21.867 ± 4.708% when treated with 240 μg/mL and 300 μg/mL concentrations of the extract, respectively. The non-apoptotic cell population increased significantly from 3.983 ± 0.631% (vehicle control) to 44.117 ± 8.818% only when treated with 300 μg/mL concentration of the extract. The viable or live cell population decreased significantly from 86.483 ± 1.311% (vehicle control) to 31.500 ± 4.906% only when treated with 300 μg/mL concentration of the extract. However, no significant change was observed in the early apoptotic cell population compared to the vehicle control. Taken together, it was observed that the late apoptotic population increased significantly in all the concentrations from 120 μg/mL to 300 μg/mL, the non-apoptotic population increased individually in all the concentrations from 60 μg/mL to 300 μg/mL, and the live cell population decreased individually in all the concentrations from 120 μg/mL to 300 μg/mL of LCLEE in the case of MDA-MB-231/pcDNA3.1(+) cells. However, in the case of MDA-MB-231/pcDNA3.1(+)-DNMT1 cells, the late apoptotic population increased significantly only at the 240 μg/mL and 300 μg/mL concentrations, and the non-apoptotic population increased significantly only at the 300 μg/mL concentration. Live cell population decreased significantly only at the 300 μg/mL concentration of LCLEE. A comparative table of live and dead cell populations at different treatment conditions is provided in Supplementary Table S5. The above findings culminated in the fact that the ectopic overexpression of DNMT1 can attenuate LCLEE-induced apoptosis.

**Figure 6.**
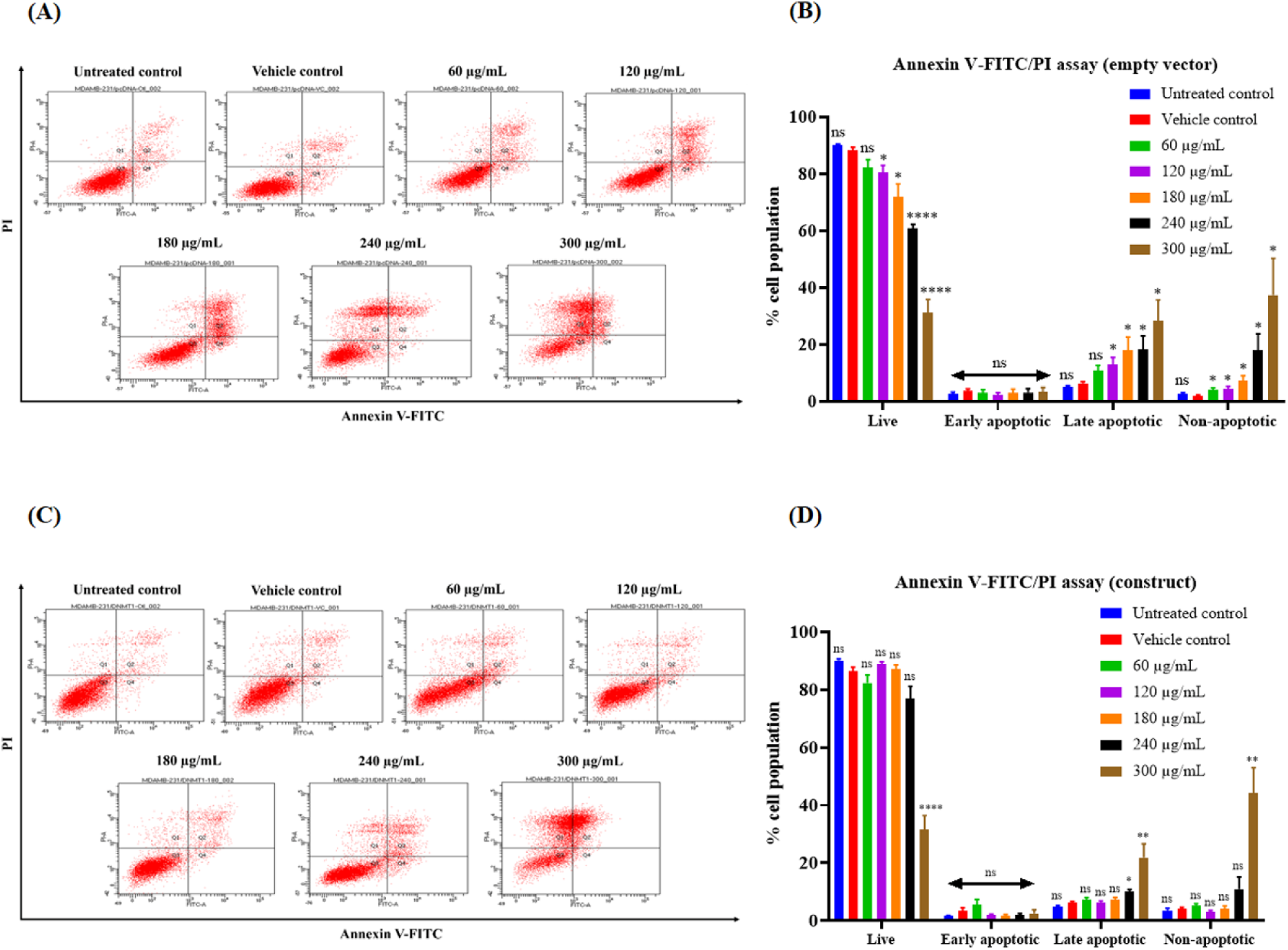
Representative scatter plots showing percentage viable or live (Q3), early apoptotic (Q4), late apoptotic (Q2), and non-apoptotic (Q1) **(A)** MDA-MB-231/pcDNA3.1(+) cells and **(C)** MDA-MB-231/pcDNA3.1(+)-DNMT1 cells following 24 hours of treatment with the extract at various doses. Bar diagrams showing the percentage of **(B)** MDA-MB-231/pcDNA3.1(+) cells and **(D)** MDA-MB-231/pcDNA3.1(+)-DNMT1 cells in different quadrants (live, early apoptotic, late apoptotic, and non-apoptotic) following 24 hours of treatment with the extract at various doses. Data are means of three independent experiments and presented as means ± SEM, where * is *P*≤0.05, ** is *P*≤0.01, **** is *P*≤0.0001, and ns signifies non-significant.

### 3.7. Ectopic overexpression of DNMT1 promoted the migratory potential

To delineate whether the downregulation of DNMT1 is responsible for the inhibition of migration by the extract, wound healing was performed in MDA-MB-231/pcDNA3.1(+) and MDA-MB-231/pcDNA3.1(+)-DNMT1 cells post-treatment with sub-IC_50_ concentrations of LCLEE, along with vehicle-treated and untreated controls. The wound closure percentage in MDA-MB-231/pcDNA3.1(+) cells was found to be 61.807 ± 2.729% for untreated control and 58.377 ± 4.630% for vehicle control, whereas the wound closure percentage reduced dose-dependently to 52.301 ± 3.732%, 44.410 ± 5.063%, 38.805 ± 3.274%, 31.763 ± 2.521%, and 22.462 ± 5.936% at 40 μg/mL, 60 μg/mL, 80 μg/mL, 100 μg/mL, and 120 μg/mL, respectively. The wound closure percentage in MDA-MB-231/pcDNA3.1(+)-DNMT1 cells was found to be 88.680 ± 4.820% for untreated control and 88.561 ± 3.707% for vehicle control, whereas the wound closure percentage became 84.829 ± 4.963%, 86.066 ± 3.723%, 79.220 ± 3.608%, 72.898 ± 3.177%, and 74.096 ± 3.345% at 40 μg/mL, 60 μg/mL, 80 μg/mL, 100 μg/mL, and 120 μg/mL, respectively. Compared to the untreated control and vehicle control, a significant impediment to wound closure percentage was observed at all concentrations of LCLEE, ranging from 60 μg/mL to 120 μg/mL in the case of MDA-MB-231/pcDNA3.1(+) cells (Figure 7A and 7B). Whereas, a significant impediment to wound closure percentage was observed only at 100 μg/mL and 120 μg/mL concentrations of LCLEE in the case of MDA-MB-231/pcDNA3.1(+)-DNMT1 cells (Figure 7C and 7D). Not only that, but when considering only the untreated cells, it was observed that the migratory potential of MDA-MB-231/pcDNA3.1(+)-DNMT1 cells is higher compared to that of MDA-MB-231/pcDNA3.1(+) cells (Figure 7E). A comparative table of wound closure percentages at various treatment conditions is provided in Supplementary Table S6. Taken together, the results indicated that the ectopic overexpression of DNMT1 promoted the migratory potential of the cells and counteracted the anti-migratory effect of LCLEE.

**Figure 7.**
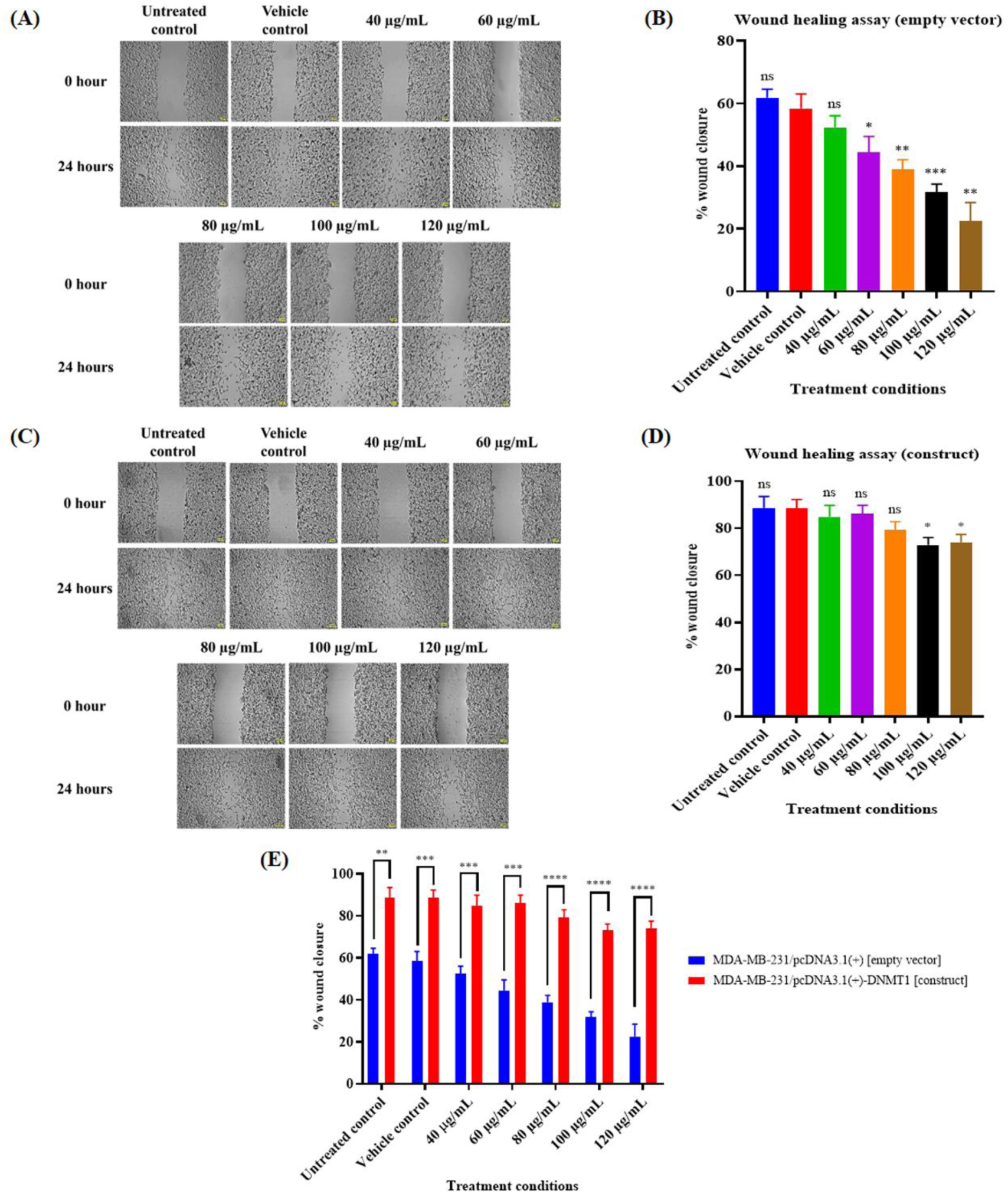
Cells were subjected to a scratch in a monolayer condition and then incubated for another 24 hours. Wound closure was visualized with a phase contrast microscope, and images were captured at 24-hour time intervals. Representative images of wound healing assay of untreated, vehicle-treated, and the extract-treated **(A)** MDA-MB-231/pcDNA3.1(+) cells and **(C)** MDA-MB-231/pcDNA3.1(+)-DNMT1 cells. Bar diagrams representing the wound closure percentage of **(B)** MDA-MB-231/pcDNA3.1(+) cells, **(D)** MDA-MB-231/pcDNA3.1(+)- DNMT1 cells, and **(E)** both the polyclonal stable cells combined at 24-hour time intervals after being treated with various doses of the extract. Data is represented as means ± SEM of three independent biological replicates, where * is *P*≤0.05, ** is *P*≤0.01, *** is *P*≤0.001, and **** is P≤0.0001.

### 3.8. DNMT1 overexpression downregulated the mRNA expression of multiple tumor suppressor genes

Previously, it was observed that LCLEE upregulated the mRNA expression of *CDKN1A*, *PTEN*, *FOXO3a*, and *Claudin-1* tumor suppressor genes (section 3.2) that remain silenced due to DNMT1-driven hypermethylation in MDA-MB-231 cells. To further check whether the ectopic overexpression of DNMT1 can alter the mRNA expression of these essential genes involved in proliferation, survival, and migration, RT-qPCR was performed, and the mRNA expression of these genes was evaluated in MDA-MB-231/pcDNA3.1(+)-DNMT1 cells compared to MDA-MB-231/pcDNA3.1(+) cells. It was found that the transcript level of *CDKN1A*, *PTEN*, *FOXO3a*, and *Claudin-1* was decreased to 0.369 ± 0.082, 0.387 ± 0.084, 0.375 ± 0.087, and 0.379 ± 0.075-fold, respectively (Figure 8) in MDA-MB-231/pcDNA3.1(+)-DNMT1 cells compared to MDA-MB-231/pcDNA3.1(+) cells.

**Figure 8.**
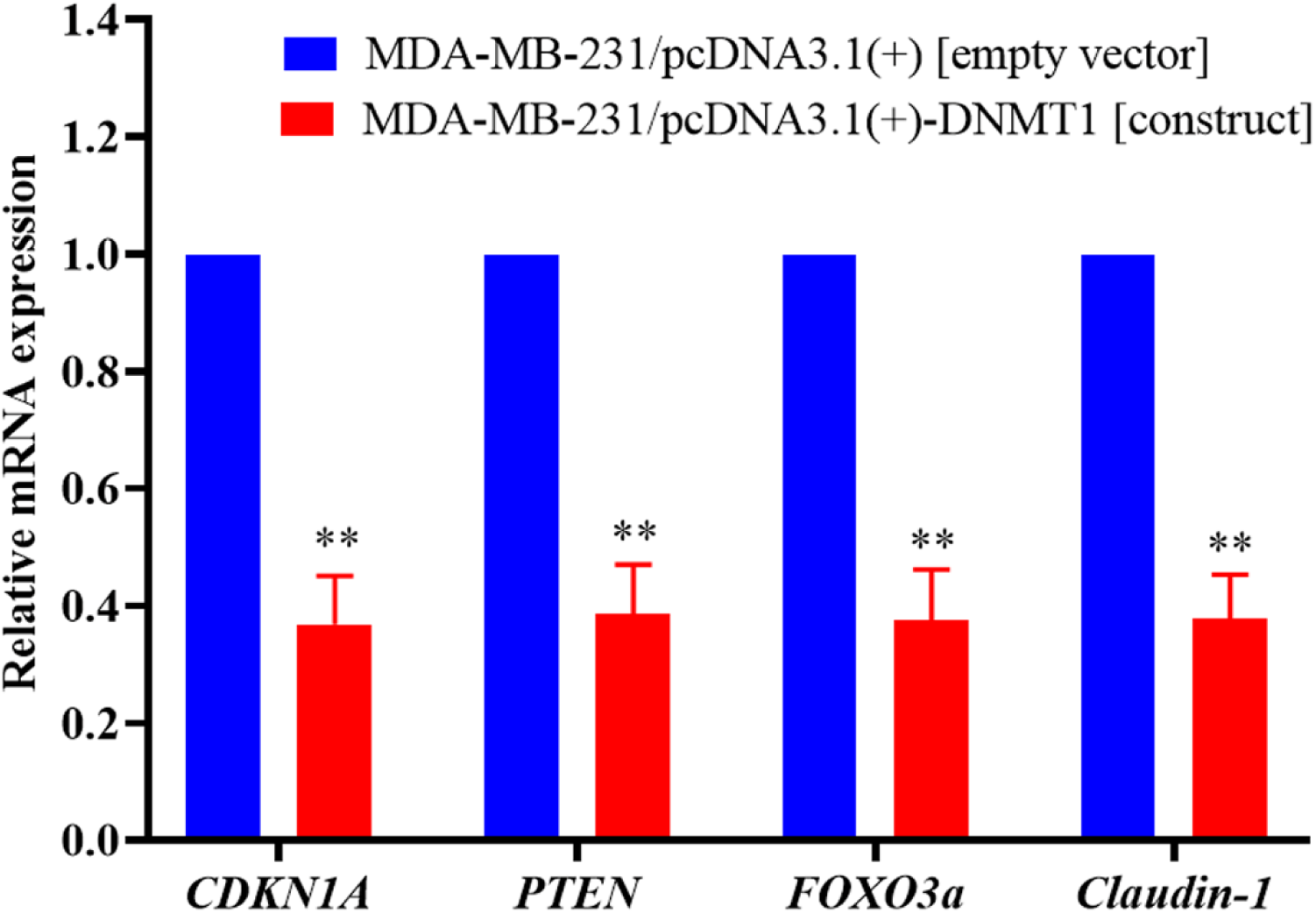
Relative mRNA expression of *CDKN1A*, *PTEN*, *FOXO3a*, and *Claudin-1* in DNMT1 overexpressed MDA-MB-231/pcDNA3.1(+)-DNMT1 cells compared to empty vector containing MDA-MB-231/pcDNA3.1(+) cells. mRNA expression was measured by RT-qPCR using specific primer pairs. The relative expressions were calculated by the 2^-ΔΔCt^ method and normalized with 18S rRNA. Data are means of three independent experiments and presented as mean ± SEM, where ** is P ≤ 0.01.

### 3.9. LCLEE downregulated the mRNA expression of *MBD2*, *HDAC1*, and *HDAC2* in MDA-MB-231 cells

The observed downregulation of DNMT1 by LCLEE suggests a potential mechanism for epigenetic reprogramming resulting in reduced maintenance of promoter hypermethylation. As a result, genes silenced via DNMT1-mediated hypermethylation are upregulated, indicating functional reversal of transcriptional repression. However, DNA methylation does not operate alone; it is tightly integrated with methyl-CpG binding proteins and histone-modifying enzymes that collectively establish and stabilize repressive chromatin states. MBD2 is a critical methyl-CpG “reader” that recognizes methylated promoters and recruits histone deacetylases, particularly HDAC1 and HDAC2, to enforce gene silencing (Aanniz et al., 2024; Lekesiz et al., 2025). Thus, even if DNMT1 levels decline, persistent expression or activity of MBD2 and HDAC1/2 may sustain transcriptional repression. Assessing the expression of MBD2, HDAC1, and HDAC2 is therefore essential to delineate the broader epigenetic impact of the extract. MBD2 mRNA expression (Figure 9A) was found to be significantly downregulated to 0.56 ± 0.11, 0.76 ± 0.05, 0.25 ± 0.12, and 0.2 ± 0.02-fold when MDA-MB-231 cells were treated with 40 µg/mL, 60 µg/mL, 120 µg/mL, and 180 µg/mL concentrations of LCLEE, respectively. The transcript level of HDAC1 (Figure 9B) was significantly decreased to 0.49 ± 0.06, 0.47 ± 0.01, 0.37 ± 0.04, and 0.29 ± 0.05-fold when MDA-MB-231 cells were treated with 40 µg/mL, 60 µg/mL, 120 µg/mL, and 180 µg/mL concentrations of LCLEE, respectively. Similarly, HDAC2 mRNA expression (Figure 9C) was found to be significantly downregulated to 0.53 ± 0.13, 0.63 ± 0.07, 0.22 ± 0.07, and 0.28 ± 0.01-fold when MDA-MB-231 cells were treated with 40 µg/mL, 60 µg/mL, 120 µg/mL, and 180 µg/mL concentrations of the extract, respectively.

**Figure 9.**
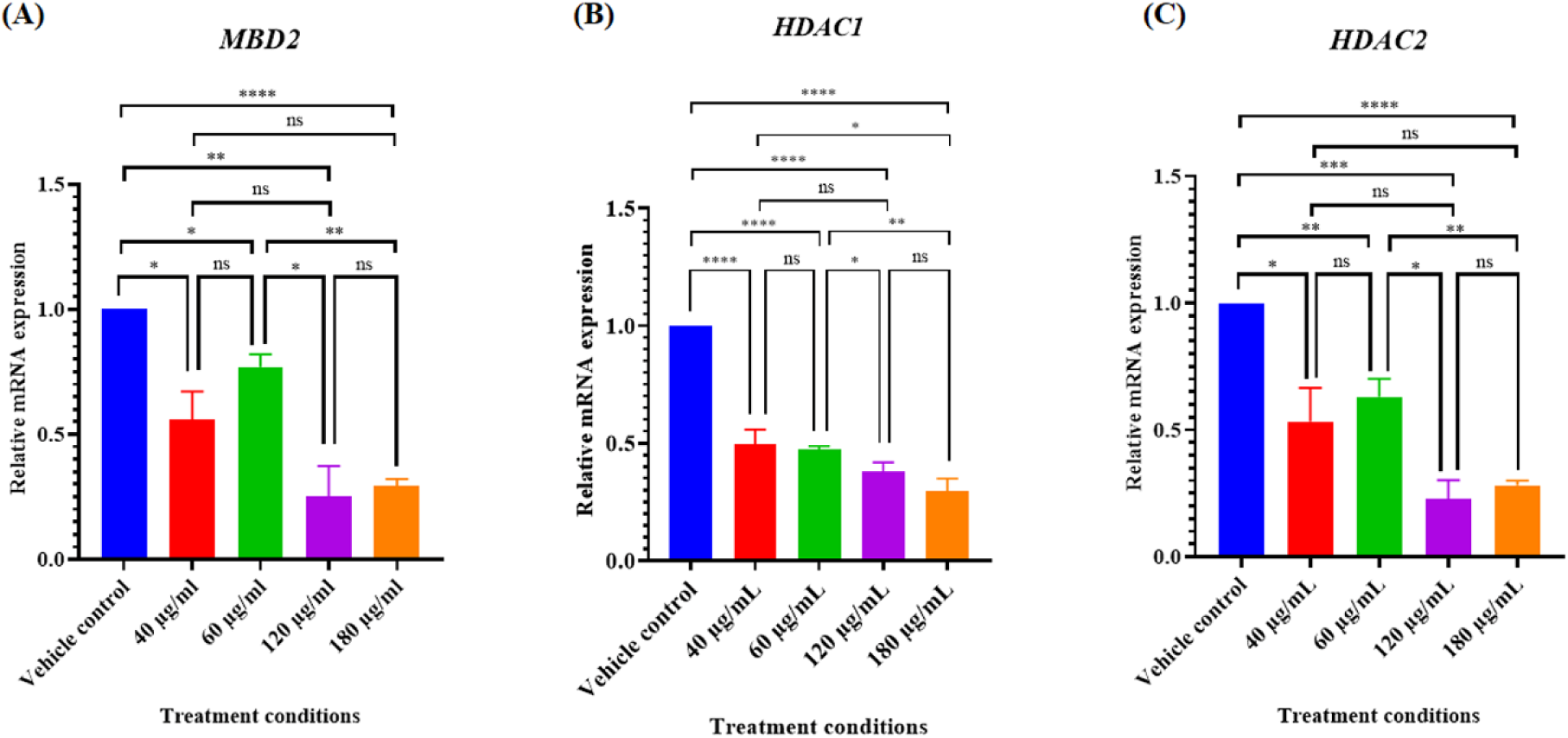
mRNA expression levels of some key epigenetic regulators were altered in MDA-MB-231 cells compared to vehicle control when treated with various doses of *Lantana camara* leaf ethanolic extract for 24 hours. Relative mRNA expression of **(A)** *MBD2*, **(B)** *HDAC1*, and **(C)** *HDAC2*. Data is represented as mean ± SEM of three independent biological replicates, where * is *P*≤0.05, ** is *P*≤0.01, *** is *P*≤0.001, **** is *P*≤0.0001, and ns signifies non-significant.

## 4. Discussion

In the present study, we found that treatment with LCLEE suppressed the expression of DNMT1 at both mRNA and protein levels in MDA-MB-231 cells (Figure 1). This suppression was accompanied by the increased expression of multiple tumor suppressor genes commonly silenced through DNMT1-driven promoter hypermethylation (Figure 2). p21, encoded by the *CDKN1A* gene, is a protein that inhibits Cyclin D-CDK4/6 complex and induces G0/G1 cell cycle arrest (Xia et al., 2023). PTEN is a crucial tumor suppressor protein that inhibits PI3K signalling by blocking PIP3-mediated membrane recruitment and activation of Akt, thereby impeding cell proliferation and survival (Miricescu et al., 2020). FOXO3a is a transcription factor belonging to the forkhead box O (FOXO) family and a tumor suppressor protein that suppresses cell cycle progression and promotes cell death (Liu et al., 2019). Claudin-1 is a tight junction protein that increases cell-cell adhesion and inhibits cell migration in TNBC cell lines (Geoffroy et al., 2020). Increased expression of these genes likely contributed to the G0/G1 arrest, apoptosis, and impaired migration observed previously (Pal et al., 2024). Importantly, ectopic overexpression of DNMT1 (Figure 3) in MDA-MB-231 cells rescued cell viability (Figure 4), impeded G0/G1 arrest (Figure 5) and apoptosis (Figure 6), promoted migration (Figure 7), and suppressed the expression of tumor suppressor genes (Figure 8), suggesting the key role of DNMT1 downregulation in mediating the anticancer activities of the extract. These findings not only support our previous report (Pal et al., 2024) but also provide mechanistic insights into the effects of the extract.

Aberrant DNA methylation, primarily mediated by DNMT1 overexpression, has been recognized as a key driver of TNBC, which lacks hormone receptors and HER2-directed therapeutic targets. Elevated DNMT1 expression has been associated with poor prognosis, tumor aggressiveness, and resistance to therapy in TNBC (Daifuku, 2025; Wong, 2021). Our observations that suppression of DNMT1 restores the expression of hypermethylated tumor suppressor genes and induces G0/G1 cell cycle arrest and apoptosis, along with the inhibition of migration, are in line with previous studies reporting DNMT1 as a crucial regulator of TNBC (H. Li et al., 2022; Z. Li et al., 2022; Pan et al., 2022).

Several studies have also reported that natural products isolated from different plants have the potential to inhibit the expression or activity of DNMT1, thereby reactivating tumor suppressor genes and mediating anticancer effects. EGCG, a primary component of green tea, increased the expression of p16 and p21 in A431 human skin cancer cells by reducing DNMT1 expression, and the activities of DNA methyltransferases and histone deacetylases (Nandakumar et al., 2011). EGCG and p-coumaric acid (found in propolis) have also been reported to reduce the viability of BT-20, BT-549, MDA-MB-231, and MDA-MB-436 triple-negative breast cancer cells. Molecular docking studies indicate that both compounds interact with the methyltransferase domain of DNMT1 and directly compete with its intrinsic inhibitor, S-adenosyl-L-homocysteine (SAH). Moreover, EGCG and the ethanolic extract of propolis partially demethylated the promoter region of RASSF1A only in the BT-549 triple-negative breast cancer cells (Assumpção et al., 2020). Brazilin, a compound found in the wood of *Caesalpinia sappan*, downregulated the expression of DNMT1 and reactivated the expression of p21 by promoter demethylation, thereby inhibiting the proliferation of MCF-7 breast cancer cells (Chatterjee et al., 2022). Kazinol Q, a phytochemical isolated from Formosan plants, has been reported to inhibit DNMT1 and to induce apoptosis in MCF-7 breast and LNCaP prostate cancer cells. The protective impact of ectopic expression of DNMT1 on Kazinol Q-induced cell death confirmed the function of DNMT1 inhibition in mediating Kazinol Q’s antiproliferative effect in MCF-7 cells (Weng et al., 2014). Curcumin, an active compound of turmeric, induced G0/G1 cell cycle arrest and apoptosis by downregulating DNMT1 and reactivating p15 tumor suppressor gene in Acute Myeloid Leukemia cell lines (Yu et al., 2013). β-elemene, a compound isolated from *Rhizoma zedoariae*, has been reported to inhibit the expression of DNMT1 and induce G0/G1 cell cycle arrest in A549 lung adenocarcinoma cells. Additionally, the effect of β-elemene on cell proliferation was reversed by the overexpression of DNMT1 (Zhao et al., 2015). Sulforaphane, a bioactive phytochemical found in cruciferous vegetables such as cabbage and broccoli, repressed the expression of DNMT1 and restored the expression of Wnt inhibitory factor 1 (WIF1), thereby reducing the cancer stem cell-associated properties and inhibiting the formation of tumor spheres in nasopharyngeal carcinoma. The siRNA knockdown studies further confirmed the role of DNMT1 (Chen et al., 2019). Genistein, a compound present in soybeans and legumes, reduced DNA methyltransferase activity and expression of DNMT1 in MCF-7 and MDA-MB-231 breast cancer cells, without any change in the expression of DNMT3A or DNMT3B. This reduction in the expression of DNMT1 resulted in the reactivation of tumor suppressor genes, including PTEN (Xie et al., 2014). A major highlight of the current work lies in the functional validation of DNMT1 suppression. The present study demonstrates that LCLEE exerts its anticancer effects in MDA-MB-231 TNBC cells through epigenetic reprogramming, specifically by suppressing the expression of DNMT1. LCLEE-mediated reactivation of tumor suppressor genes and impairment of proliferation, survival, and migration address three critical hallmarks of cancer. Given the lack of effective targeted therapies for TNBC, these results highlight the potential of phytochemicals present in the extract as novel epigenetic modulators. The identification of bioactive compound/s responsible for the epigenetic modulatory and anticancer effects of the extract, along with *in vivo* studies and genome-wide methylation profiling, will further enrich the findings of the present study. Moreover, the ability of the extract to simultaneously inhibit proliferation, survival, and migration by inhibiting DNMT1 strengthens its relevance as an epigenetic therapeutic agent for further preclinical development.

In addition to DNMT1, LCLEE also downregulated the mRNA expression of *MBD2*, *HDAC1*, and *HDAC2* (Figure 9) in MDA-MB-231 cells. These genes are well-known epigenetic corepressors that cooperate to form transcriptional silencing complexes, maintaining stable repression of tumor suppressor genes. Previous studies have reported that MBD2 and HDAC1/2 work in tandem with DNA methylation to silence tumor suppressor genes, thereby contributing to tumorigenesis (Aanniz et al., 2024; Lekesiz et al., 2025). Not only that, but MBD2 also helps in breast cancer progression, and deletion of MBD2 inhibits tumor growth by impairing the PI3K/Akt pathway and metastasis by altering the expression of N-cadherin and osteopontin *in vivo* (Mahmood et al., 2024). 5-aza-2ʹ-deoxycytidine is a nucleoside analogue that inhibits DNMTs and impedes the proliferation of non-invasive breast cancer cells. However, it paradoxically enhances the invasiveness by inducing pro-metastatic gene expression. On the other hand, MBD2 silences methylated tumor suppressor genes and activates pro-metastatic genes. Cheishvili and colleagues suggested that dual targeting of MBD2 and DNMTs is a more effective epigenetic therapeutic strategy. The combined depletion of MBD2 and inhibition of DNMTs induces G0/G1 arrest and apoptosis, while simultaneously counteracting the invasiveness induced by DNMT inhibition alone by 5-aza-2ʹ-deoxycytidine, both *in vitro* and *in vivo* (Cheishvili et al., 2014). EGCG also suppressed the expression of DNMT1, HDAC1, and HDAC2, and inhibited the proliferation and survival of NB4 and HL-60 acute promyelocytic leukemia cells by inducing G0/G1 cell cycle arrest and apoptosis (Borutinskaitė et al., 2018). Moreover, C02S, a dual inhibitor of DNMT and HDAC, inhibited migration, invasion, and angiogenesis, induced G0/G1 cell cycle arrest and apoptosis, and suppressed tumour growth in breast cancer, both *in vitro* and *in vivo* (Yuan et al., 2019). The aforementioned studies support the findings of the present study, indicating that the simultaneous downregulation of DNMT1, MBD2, HDAC1, and HDAC2 by LCLEE exerts a synergistic effect, leading to the induction of G0/G1 cell cycle arrest and apoptosis, and inhibition of migration, thereby enhancing the anticancer efficacy of the extract.

Previous studies have reported that *Lantana camara* leaf extract and derived compounds show antiproliferative and cytotoxic activities against various cancer types, including breast cancer (El-Din et al., 2022; Han et al., 2015; Shamsee et al., 2019). However, apart from our earlier report (Pal et al., 2024), there are no detailed studies that elucidate the effect of *Lantana camara* extract in TNBC. In continuation of that, the present study is the first of its kind to provide a strong mechanistic insight, unravelling that the downregulation of DNMT1 is not merely an associated event but is mechanistically central to the anticancer activity of *Lantana camara* leaf extract.

## 5. Conclusion

In summary, *Lantana camara* leaf ethanolic extract downregulates DNMT1 and associated epigenetic repressors, leading to the re-expression of silenced tumor suppressor genes, the promotion of G0/G1 cell cycle arrest and apoptosis, and the suppression of migration in MDA-MB-231 triple-negative breast cancer cells. Ectopic overexpression studies further confirmed the mechanistic link between DNMT1 suppression and these observed changes. These findings not only provide mechanistic insights into the anticancer activity of the extract but also highlight its potential as a natural epigenetic therapeutic candidate for triple-negative breast cancer.

## Acknowledgement

SS, AP, and TKS acknowledge the Indian Institute of Science Education and Research Kolkata (IISER Kolkata) for the research facilities. SS and AP recognize the Department of Biotechnology (DBT), Government of India, for research fellowships. The authors would like to acknowledge Prof. Subhajit Bandyopadhyay, Department of Chemical Sciences, IISER Kolkata, for providing the rotary evaporator facility. The authors would also like to convey their gratitude to Dr. Sumita Sengupta (nee Bandyopadhyay), Department of Biophysics, Molecular Biology and Bioinformatics, University of Calcutta, for providing the pcDNA3.1(+) mammalian expression vector. The authors would also like to acknowledge Mr. Tamal Ghosh for his technical support in flow cytometry.

## Authors’ contributions

**Sourav Sanyal:** Conceptualization, Methodology, Investigation, Formal analysis, Writing – original draft, Writing – Review & Editing. **Arundhaty Pal:** Methodology, Investigation, Formal analysis, Writing – original draft, Writing – Review & Editing. **Tapas K Sengupta:** Conceptualization, Writing – original draft, Writing – Review & Editing, Supervision, Project administration, Funding acquisition.

## Data Availability

All data generated or analysed during this study are included in this present article and supplementary section.

## Declaration

### Ethical approval

Not applicable.

### Consent to participate

All authors have given consent to participate in the study.

### Consent for publication

All authors have given consent for publication.

## References

Aanniz, T., Bouyahya, A., Balahbib, A., El Kadri, K., Khalid, A., Makeen, H.A., Alhazmi, H.A., El Omari, N., Zaid, Y., Wong, R.S.Y., Yeo, C.I., Goh, B.H., Bakrim, S., 2024. Natural bioactive compounds targeting DNA methyltransferase enzymes in cancer: Mechanisms insights and efficiencies. Chem Biol Interact 392, 110907. 10.1016/J.CBI.2024.110907

Al-Kharashi, L.A., Al-Mohanna, F.H., Tulbah, A., Aboussekhra, A., 2017. The DNA methyl-transferase protein DNMT1 enhances tumor-promoting properties of breast stromal fibroblasts. Oncotarget 9, 2329. 10.18632/ONCOTARGET.23411

Assumpção, J.H.M., Takeda, A.A.S., Sforcin, J.M., Rainho, C.A., 2020. Effects of Propolis and Phenolic Acids on Triple-Negative Breast Cancer Cell Lines: Potential Involvement of Epigenetic Mechanisms. Molecules 25, 1289. 10.3390/MOLECULES25061289

Borutinskaitė, V., Virkšaitė, A., Gudelytė, G., Navakauskienė, R., 2018. Green tea polyphenol EGCG causes anti-cancerous epigenetic modulations in acute promyelocytic leukemia cells. Leuk Lymphoma 59, 469–478. 10.1080/10428194.2017.1339881

Bray, F., Laversanne, M., Sung, H., Ferlay, J., Siegel, R.L., Soerjomataram, I., Jemal, A., 2024. Global cancer statistics 2022: GLOBOCAN estimates of incidence and mortality worldwide for 36 cancers in 185 countries. CA Cancer J Clin 74, 229–263. 10.3322/CAAC.21834

Chatterjee, B., Ghosh, K., Swain, A., Nalla, K.K., Ravula, H., Pan, A., Kanade, S.R., 2022. The phytochemical brazilin suppress DNMT1 expression by recruiting p53 to its promoter resulting in the epigenetic restoration of p21 in MCF7cells. Phytomedicine 95, 153885. 10.1016/J.PHYMED.2021.153885

Cheishvili, D., Chik, F., Li, C.C. hen, Bhattacharya, B., Suderman, M., Arakelian, A., Hallett, M., Rabbani, S.A., Szyf, M., 2014. Synergistic effects of combined DNA methyltransferase inhibition and MBD2 depletion on breast cancer cells; MBD2 depletion blocks 5-aza-2ʹ-deoxycytidine-triggered invasiveness. Carcinogenesis 35, 2436. 10.1093/CARCIN/BGU181

Chen, L., Chan, L.S., Lung, H.L., Yip, T.T.C., Ngan, R.K.C., Wong, J.W.C., Lo, K.W., Ng, W.T., Lee, A.W.M., Tsao, G.S.W., Lung, M.L., Mak, N.K., 2019. Crucifera sulforaphane (SFN) inhibits the growth of nasopharyngeal carcinoma through DNA methyltransferase 1 (DNMT1)/Wnt inhibitory factor 1 (WIF1) axis. Phytomedicine 63, 153058. 10.1016/J.PHYMED.2019.153058

Chiang, S.K., Chang, W.C., Chen, S.E., Chang, L.C., 2019. DOCK1 Regulates Growth and Motility through the RRP1B-Claudin-1 Pathway in Claudin-Low Breast Cancer Cells. Cancers (Basel) 11, 1762. 10.3390/CANCERS11111762

Daifuku, R., 2025. The promise of DNA methyltransferase inhibitors for the treatment of triple negative breast cancer. Med Res Arch 13. 10.18103/MRA.V13I2.6276

El-Din, M.I.G., Fahmy, N.M., Wu, F., Salem, M.M., Khattab, O.M., El-Seedi, H.R., Korinek, M., Hwang, T.L., Osman, A.K., El-Shazly, M., Fayez, S., 2022. Comparative LC-LTQ-MS-MS Analysis of the Leaf Extracts of Lantana camara and Lantana montevidensis Growing in Egypt with Insights into Their Antioxidant, Anti-Inflammatory, and Cytotoxic Activities. Plants (Basel) 11. 10.3390/PLANTS11131699

Geoffroy, M., Kleinclauss, A., Kuntz, S., Grillier-Vuissoz, I., 2020. Claudin 1 inhibits cell migration and increases intercellular adhesion in triple-negative breast cancer cell line. Mol Biol Rep 47, 7643–7653. 10.1007/S11033-020-05835-3/FIGURES/4

Han, E.B., Chang, B.Y., Jung, Y.S., Kim, S.Y., 2015. Lantana camara Induces Apoptosis by Bcl-2 Family and Caspases Activation. Pathol Oncol Res 21, 325–331. 10.1007/S12253-014-9824-4

Lee, J., 2023. Current Treatment Landscape for Early Triple-Negative Breast Cancer (TNBC). J Clin Med 12, 1524. 10.3390/JCM12041524

Lekesiz, R.T., Koca, K.K., Kugu, G., Çalışkaner, Z.O., 2025. Versatile functions of methyl-CpG-binding domain 2 (MBD2) in cellular characteristics and differentiation. Mol Biol Rep 52, 1–12. 10.1007/S11033-025-10411-8/FIGURES/1

Li, H., Li, H.H., Chen, Q., Wang, Y.Y., Fan, C.C., Duan, Y.Y., Huang, Y., Zhang, H.M., Li, J.P., Zhang, X.Y., Xiang, Y., Gu, C.J., Wang, L., Liao, X.H., Zhang, T.C., 2022. miR-142-5p Inhibits Cell Invasion and Migration by Targeting DNMT1 in Breast Cancer. Oncol Res 28, 885. 10.3727/096504021X16274672547967

Li, Z., Wang, P., Cui, W., Yong, H., Wang, D., Zhao, T., Wang, W., Shi, M., Zheng, J., Bai, J., 2022. Tumour-associated macrophages enhance breast cancer malignancy via inducing ZEB1-mediated DNMT1 transcriptional activation. Cell Biosci 12, 176. 10.1186/S13578-022-00913-4

Liang, F., Zhang, H., Gao, H., Cheng, D., Zhang, N., Du, J., Yue, J., Du, P., Zhao, B., Yin, L., 2020. Liquiritigenin decreases tumorigenesis by inhibiting DNMT activity and increasing BRCA1 transcriptional activity in triple-negative breast cancer. Exp Biol Med 246, 459. 10.1177/1535370220957255

Liu, H., Song, Y., Qiu, H., Liu, Y., Luo, K., Yi, Y., Jiang, G., Lu, M., Zhang, Z., Yin, J., Zeng, S., Chen, X., Deng, M., Jia, X., Gu, Y., Chen, D., Zheng, G., He, Z., 2019. Downregulation of FOXO3a by DNMT1 promotes breast cancer stem cell properties and tumorigenesis. Cell Death Differ 27, 966. 10.1038/S41418-019-0389-3

Liu, T., Wang, J., Sun, L., Li, M., He, X., Jiang, J., Zhou, Q., 2021. Piwi-interacting RNA-651 promotes cell proliferation and migration and inhibits apoptosis in breast cancer by facilitating DNMT1-mediated PTEN promoter methylation. Cell Cycle 20, 1603. 10.1080/15384101.2021.1956090

Livak, K.J., Schmittgen, T.D., 2001. Analysis of Relative Gene Expression Data Using Real-Time Quantitative PCR and the 2−ΔΔCT Method. Methods 25, 402–408. 10.1006/METH.2001.1262

Lu, Y., Chan, Y.T., Tan, H.Y., Li, S., Wang, N., Feng, Y., 2020. Epigenetic regulation in human cancer: the potential role of epi-drug in cancer therapy. Mol Cancer 19, 79. 10.1186/S12943-020-01197-3

Mahmood, N., Arakelian, A., Szyf, M., Rabbani, S.A., 2024. Methyl-CpG binding domain protein 2 (Mbd2) drives breast cancer progression through the modulation of epithelial-to-mesenchymal transition. Experimental & Molecular Medicine 2024 56:4 56, 959–974. 10.1038/s12276-024-01205-2

Miricescu, D., Totan, A., Stanescu-Spinu, I.I., Badoiu, S.C., Stefani, C., Greabu, M., 2020. PI3K/AKT/mTOR Signaling Pathway in Breast Cancer: From Molecular Landscape to Clinical Aspects. Int J Mol Sci 22, 173. 10.3390/IJMS22010173

Mirza, S., Sharma, G., Parshad, R., Gupta, S.D., Pandya, P., Ralhan, R., 2013. Expression of DNA Methyltransferases in Breast Cancer Patients and to Analyze the Effect of Natural Compounds on DNA Methyltransferases and Associated Proteins. J Breast Cancer 16, 23. 10.4048/JBC.2013.16.1.23

Nandakumar, V., Vaid, M., Katiyar, S.K., 2011. (−)-Epigallocatechin-3-gallate reactivates silenced tumor suppressor genes, Cip1/p21 and p16INK4a, by reducing DNA methylation and increasing histones acetylation in human skin cancer cells. Carcinogenesis 32, 537. 10.1093/CARCIN/BGQ285

Pal, A., Sanyal, S., Das, S., Sengupta, T.K., 2024. Effect of Lantana camara Ethanolic Leaf Extract on Survival and Migration of MDA-MB-231 Triple-Negative Breast Cancer Cell Line. J Herb Med 43, 100837. 10.1016/J.HERMED.2023.100837

Pan, T., Ding, H., Jin, L., Zhang, S., Wu, D., Pan, W., Dong, M., Ma, X., Chen, Z., 2022. DNMT1-mediated demethylation of lncRNA MEG3 promoter suppressed breast cancer progression by repressing Notch1 signaling pathway. Cell Cycle 21, 2323. 10.1080/15384101.2022.2094662

Saldívar-González, F.I., Gómez-García, A., Chávez-Ponce De León, D.E., Sánchez-Cruz, N., Ruiz-Rios, J., Pilón-Jiménez, B.A., Medina-Franco, J.L., 2018. Inhibitors of DNA Methyltransferases From Natural Sources: A Computational Perspective. Front Pharmacol 9, 1144. 10.3389/FPHAR.2018.01144

Shamsee, Z.R., Al-Saffar, A.Z., Al-Shanon, A.F., Al-Obaidi, J.R., 2019. Cytotoxic and cell cycle arrest induction of pentacyclic triterpenoides separated from Lantana camara leaves against MCF-7 cell line in vitro. Mol Biol Rep 46, 381–390. 10.1007/S11033-018-4482-3

Soumya, T., Lakshmipriya, T., Klika, K.D., Jayasree, P.R., Manish Kumar, P.R., 2021. Anticancer potential of rhizome extract and a labdane diterpenoid from Curcuma mutabilis plant endemic to Western Ghats of India. Sci Rep 11, 552. 10.1038/S41598-020-79414-8

Stefanska, B., Salamé, P., Bednarek, A., Fabianowska-Majewska, K., 2012. Comparative effects of retinoic acid, vitamin D and resveratrol alone and in combination with adenosine analogues on methylation and expression of phosphatase and tensin homologue tumour suppressor gene in breast cancer cells. British Journal of Nutrition 107, 781–790. 10.1017/S0007114511003631

Weng, J.R., Lai, I.L., Yang, H.C., Lin, C.N., Bai, L.Y., 2014. Identification of Kazinol Q, a Natural Product from Formosan Plants, as an Inhibitor of DNA Methyltransferase. Phytotherapy Research 28, 49–54. 10.1002/PTR.4955

Wong, K.K., 2021. DNMT1: A key drug target in triple-negative breast cancer. Semin Cancer Biol 72, 198–213. 10.1016/J.SEMCANCER.2020.05.010

Xia, W., Gong, E. sheng, Lin, Y., Zheng, B., Yang, W., Li, T., Zhang, S., Li, P., Liu, R. hai, 2023. Wild pink bayberry free phenolic extract induces mitochondria-dependent apoptosis and G0/G1 cell cycle arrest through p38/MAPK and PI3K/Akt pathway in MDA-MB-231 cancer cells. Food Science and Human Wellness 12, 1510–1518. 10.1016/J.FSHW.2023.02.014

Xie, Q., Bai, Q., Zou, L.Y., Zhang, Q.Y., Zhou, Y., Chang, H., Yi, L., Zhu, J.D., Mi, M.T., 2014. Genistein inhibits DNA methylation and increases expression of tumor suppressor genes in human breast cancer cells. Genes Chromosomes Cancer 53, 422–431. 10.1002/GCC.22154

Yin, L., Duan, J.J., Bian, X.W., Yu, S.C., 2020. Triple-negative breast cancer molecular subtyping and treatment progress. Breast Cancer Res 22, 61. 10.1186/S13058-020-01296-5

Yu, J., Peng, Y., Wu, L.C., Xie, Z., Deng, Y., Hughes, T., He, S., Mo, X.K., Chiu, M., Wang, Q.E., He, X., Liu, S., Grever, M.R., Chan, K.K., Liu, Z., 2013. Curcumin Down-Regulates DNA Methyltransferase 1 and Plays an Anti-Leukemic Role in Acute Myeloid Leukemia. PLoS One 8, e55934. 10.1371/JOURNAL.PONE.0055934

Yu, J., Zayas, J., Qin, B., Wang, L., 2019. Targeting DNA methylation for treating triple-negative breast cancer. Pharmacogenomics 20, 1151. 10.2217/PGS-2019-0078

Yuan, Z., Chen, S., Gao, C., Dai, Q., Zhang, C., Sun, Q., Lin, J.S., Guo, C., Chen, Y., Jiang, Y., 2019. Development of a versatile DNMT and HDAC inhibitor C02S modulating multiple cancer hallmarks for breast cancer therapy. Bioorg Chem 87, 200–208. 10.1016/J.BIOORG.2019.03.027

Zhao, S.Y., Wu, J., Zheng, F., Tang, Q., Yang, L.J., Li, L., Wu, W.Y., Hann, S.S., 2015. β-elemene inhibited expression of DNA methyltransferase 1 through activation of ERK1/2 and AMPKα signalling pathways in human lung cancer cells: the role of Sp1. J Cell Mol Med 19, 630. 10.1111/JCMM.12476

Zhu, L., Tang, N., Hang, H., Zhou, Y., Dong, J., Yang, Y., Mao, L., Qiu, Y., Fu, X., Cao, W., 2024. Loss of Claudin-1 incurred by DNMT aberration promotes pancreatic cancer progression. Cancer Lett 586. 10.1016/j.canlet.2024.216611

