## Supplementary File for "*Lantana camara* leaf extract-mediated suppression of DNA methyltransferase 1 promotes G0/G1 cell cycle arrest and apoptosis, and impedes migration in MDA-MB-231 triple-negative breast cancer cells"

**Table S1.** Details of primers used in the RT-qPCR experiment.

| Gene | Forward Primer (5´-3´) | Reverse Primer (5´-3´) | Amplicon Size (bp) |
|---|---|---|---|
| ***18S*** | ACGGAAGGGCACCACCAGGAGT | GAACGGCCATGCACCACCACC | 138 |
| ***DNMT1*** | TGCCAAACGGAGGCCCGAAGA | CTTGGGAGGGTGGGTCTTGGAG | 175 |
| ***CDKN1A*** | CACTCAGAGGAGGCGCCATGTC | ATCGCTCACGGGCCTCCTGGAT | 162 |
| ***PTEN*** | TGGCGGAACTTGCAATCCTCAGT | ACCACACACAGGTAACGGCTGAG | 133 |
| ***FOXO3a*** | TTGCAGAACTCCATCCGGCACA | GCGGCCACGGCTCTTGGTAT | 183 |
| ***Claudin-1*** | GTGCTTGGAAGACGATGAGGTGC | CGTACCTGGCATTGACTGGGGTC | 161 |
| ***MBD2*** | GCTAAGTGCTGGCAAGAGCGA | GAGAGGATCGTTTCGCAGTCTCTG | 184 |
| ***HDAC1*** | CTGGGGACCTACGGGATATCGGG | CCCCAGATAGGGAGTCTGAGCCA | 183 |
| ***HDAC2*** | CCCATAAAGCCACTGCCGAAGAA | ACAGCTCCAGCAACTGAACCG | 199 |

**Table S2.** Details of the antibodies used in the present study.

| Antibody | Source | Manufacturer | Catalogue No. | Dilution used | MW (kDa) |
|---|---|---|---|---|---|
| Anti-GAPDH | Rabbit | Affinity Biosciences, Cincinnati, Ohio, USA | AF7021 | 1:3000 | 37 |
| Anti-DNMT1 | Rabbit | Affinity Biosciences, Cincinnati, Ohio, USA | DF7376 | 1:1000 | 183 |
| Anti-rabbit IgG, HRP-linked (Secondary) | Goat | Cell Signaling Technology, Danvers, Massachusetts, USA | 7074 | 1:5000 | - |

### Molecular cloning of DNMT1 coding region into pcDNA3.1(+) vector

For ectopic overexpression of DNMT1 in MDA-MB-231 cells, the human DNMT1 coding region (NCBI Reference sequence: NM_001379.4) was amplified by PCR amplification of cDNA synthesized from total RNA of MDA-MB-231 cells using forward primer containing a leader sequence, KpnI restriction site, Kozak sequence, and sequence specific for DNMT1 coding region including start codon and reverse primer containing sequence specific for DNMT1 coding region including stop codon, EcoRI restriction site, and a leader sequence (primer sequences used in PCR amplification for cloning are provided in Table S3). The amplified product was cloned into the KpnI-EcoRI sites of pcDNA3.1(+) mammalian expression vector (a generous gift from Dr. Sumita Sengupta (nee Bandyopadhyay), Department of Biophysics, Molecular Biology and Bioinformatics, University of Calcutta). The map of the parent vector, i.e., pcDNA3.1(+), and the map of the pcDNA3.1(+)-DNMT1 construct are illustrated in Figure S1. The maps were designed using SnapGene software (Dotmatics, Boston, Massachusetts, USA). Therefore, the multiple cloning site region between the KpnI and EcoRI sites was replaced by a DNA segment containing a Kozak consensus sequence, an ATG start codon, the DNMT1 coding region, and a stop codon. The resulting construct was designated as pcDNA3.1(+)-DNMT1, and its identity was confirmed by DNA sequencing.

**Table S3.** Primer sequences used in PCR amplification for cloning.

| Name | Forward Primer (5´-3´) | Reverse Primer (5´-3´) |
|---|---|---|
| ***DNMT1* coding region** | TAAGCAGGTACCGCCATGCCGGCGCGTACCGC | TGCTTAGAATTCCTAGTCCTTAGCAGCTTCCTCCTCC |

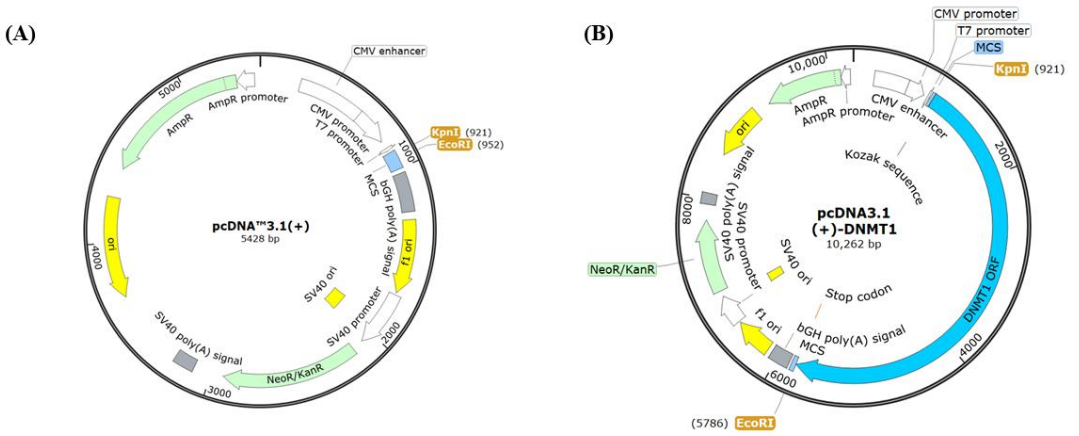


**Figure S1.** Maps of (A) pcDNA3.1(+) empty vector and (B) pcDNA3.1(+)-DNMT1 construct.

**Table S4.** Comparative analysis of cell cycle phase distribution of MDA-MB-231/pcDNA3.1(+) cells and MDA-MB-231/pcDNA3.1(+)-DNMT1 cells at different treatment conditions.

| **Treatment conditions** | **MDA-MB-231/pcDNA3.1(+) [Empty vector]** | | | **MDA-MB-231/pcDNA3.1(+)-DNMT1 [Construct]** | | |
|---|---|---|---|---|---|---|
| | **G0/G1** | **S** | **G2/M** | **G0/G1** | **S** | **G2/M** |
| **Untreated control** | 60.033 ± 0.691 | 18.783 ± 0.35 | 21.00 ± 0.831 | 66.683 ± 1.637 | 13.617 ± 0.254 | 18.917 ± 1.406 |
| **Vehicle control** | 60.967 ± 0.878 | 18.133 ± 0.481 | 20.650 ± 1.135 | 66.917 ± 1.097 | 12.750 ± 0.348 | 19.717 ± 0.804 |
| **60 µg/mL LCLEE** | 65.917 ± 1.049 | 15.533 ± 1.067 | 17.467 ± 0.520 | 69.133 ± 0.304 | 10.517 ± 0.189 | 19.500 ± 0.434 |
| **80 µg/mL LCLEE** | 68.833 ± 1.160 | 13.550 ± 0.732 | 16.717 ± 0.611 | 71.083 ± 0.776 | 8.950 ± 0.362 | 19.083 ± 0.709 |
| **100 µg/mL LCLEE** | 72.467 ± 0.495 | 11.417 ± 0.310 | 15.300 ± 0.523 | 72.617 ± 0.516 | 7.250 ± 0.177 | 19.217 ± 0.389 |
| **120 µg/mL LCLEE** | 74.517 ± 0.588 | 9.983 ± 0.383 | 14.600 ± 0.500 | 73.400 ± 0.842 | 6.800 ± 0.267 | 18.833 ± 0.764 |
| **150 µg/mL LCLEE** | 72.817 ± 1.131 | 9.467 ± 0.189 | 16.850 ± 0.839 | 73.133 ± 0.638 | 6.083 ± 0.204 | 19.283 ± 0.597 |
| **180 µg/mL LCLEE** | 68.700 ± 1.479 | 10.800 ± 0.353 | 19.450 ± 0.880 | 69.167 ± 0.429 | 7.350 ± 0.451 | 21.850 ± 0.533 |

**Table S5.** Comparative analysis of live and dead cell populations of MDA-MB-231/pcDNA3.1(+) cells and MDA-MB-231/pcDNA3.1(+)-DNMT1 cells at different treatment conditions.

| **Treatment conditions** | **MDA-MB-231/pcDNA3.1(+) [Empty vector]** | | | | **MDA-MB-231/pcDNA3.1(+)-DNMT1 [Construct]** | | | |
|---|---|---|---|---|---|---|---|---|
| | **Live** | **Early apoptotic** | **Late apoptotic** | **Non-apoptotic** | **Live** | **Early apoptotic** | **Late apoptotic** | **Non-apoptotic** |
| **Untreated control** | 89.925 ± 0.502 | 2.450 ± 0.756 | 5.025 ± 0.475 | 2.575 ± 0.477 | 90.067 ± 0.544 | 1.633 ± 0.214 | 4.717 ± 0.527 | 3.600 ± 0.679 |
| **Vehicle control** | 88.250 ± 1.068 | 3.625 ± 0.743 | 6.275 ± 0.557 | 1.900 ± 0.319 | 86.483 ± 1.311 | 3.350 ± 1.061 | 6.200 ± 0.484 | 3.983 ± 0.631 |
| **60 µg/mL LCLEE** | 82.300 ± 2.664 | 2.950 ± 1.117 | 10.700 ± 1.901 | 4.075 ± 0.680 | 82.333 ± 2.743 | 5.467 ± 1.923 | 7.150 ± 0.855 | 5.067 ± 0.774 |
| **120 µg/mL LCLEE** | 80.350 ± 2.547 | 2.175 ± 0.832 | 12.975 ± 2.478 | 4.475 ± 0.829 | 88.750 ± 0.841 | 1.883 ± 0.421 | 6.300 ± 0.534 | 3.067 ± 0.515 |
| **180 µg/mL LCLEE** | 71.925 ± 4.486 | 2.975 ± 1.314 | 17.850 ± 4.777 | 7.275 ± 1.736 | 87.033 ± 1.536 | 1.700 ± 0.459 | 7.267 ± 0.759 | 4.017 ± 1.093 |
| **240 µg/mL LCLEE** | 60.700 ± 1.512 | 3.075 ± 1.349 | 18.175 ± 4.826 | 18.025 ± 5.693 | 76.933 ± 4.130 | 2.000 ± 0.499 | 10.083 ± 0.712 | 10.967 ± 4.100 |
| **300 µg/mL LCLEE** | 31.175 ± 4.669 | 3.175 ± 1.713 | 28.275 ± 7.393 | 37.400 ± 12.906 | 31.500 ± 4.906 | 2.533 ± 1.226 | 21.867 ± 4.708 | 44.117 ± 8.818 |

**Table S6.** Comparative analysis of the wound closure percentages in MDA-MB-231/pcDNA3.1(+) cells and MDA-MB-231/pcDNA3.1(+)-DNMT1 cells at different treatment conditions after 24 hours of treatment.

| **Treatment conditions** | **MDA-MB-231/pcDNA3.1(+) [Empty vector]** | **MDA-MB-231/pcDNA3.1(+)-DNMT1 [Construct]** |
|---|---|---|
| **Untreated control** | 61.807 ± 2.729 | 88.680 ± 4.820 |
| **Vehicle control** | 58.377 ± 4.630 | 88.561 ± 3.707 |
| **40 µg/mL LCLEE** | 52.301 ± 3.732 | 84.829 ± 4.963 |
| **60 µg/mL LCLEE** | 44.410 ± 5.063 | 86.066 ± 3.723 |
| **80 µg/mL LCLEE** | 38.805 ± 3.274 | 79.220 ± 3.608 |
| **100 µg/mL LCLEE** | 31.763 ± 2.521 | 72.898 ± 3.177 |
| **120 µg/mL LCLEE** | 22.462 ± 5.936 | 74.096 ± 3.345 |
